# An oncolytic herpesvirus expressing a CXCR4 antagonist interferes with Glioblastoma cells stemness features and migration

**DOI:** 10.1101/2025.05.21.655102

**Authors:** D’arrigo Paolo, Dubois Maxime, Sanchez Gil Judit, Lassence Cédric, Brouwers Benoit, Lombard Arnaud, Rogister Bernard, Neirinckx Virginie, Lebrun Marielle, Sadzot-Delvaux Catherine

## Abstract

Glioblastoma is one of the most aggressive brain tumors. Despite the standard therapy, the survival from diagnosis remains dramatically low, especially due to relapses. Considering their capacity to escape the tumor and to migrate to the subventricular zone through a CXCR4-dependant mechanism, Glioblastoma stem-like cells (GSCs) are considered as responsible for these relapses. CXCR4 regulates biological features associated with tumor progression, including self-renewal, migration and radio-resistance. Importantly, its expression correlates with severity and poor prognosis of several cancers including GBM. CXCR4/CXCL12 pathway appears therefore as an interesting potential therapeutic target. We have generated an oncolytic herpesvirus (oHSV) expressing HA-P2G, a mutated form of CXCL12 previously described as a CXCR4 competitive inhibitor. We demonstrate that *in vitro*, oHSV/P2G impairs human primary GSCs’ stemness markers expression, self-renewal and migration. In an orthotopic xenograft murine model, its injection within the tumor limits tumor growth and GSCs migration. The ability of P2G to interfere with major GSC features demonstrates the interest in considering oHSV/P2G as a promising new therapeutic approach for glioblastoma patients.

## INTRODUCTION

Glioblastoma (GBM) is one of the most malignant brain tumors. Despite aggressive standard therapy, consisting in surgical resection followed by radio and chemotherapy, the current overall survival is only 15-16 months from the diagnosis ^1^, due to systematic recurrence. Recurrent tumors usually display poor sensitivity to standard therapies and high infiltrative capacity ^2^.

Several studies have identified Glioblastoma stem-like cells (GSCs) as partially responsible for GBM recurrence, and aggressiveness ^3^. Historically identified for their self-renewal abilities and the expression of neural stem cell markers, GSCs have also been linked with increased invasive properties. In an orthotopic xenograft murine model GSCs have been shown to escape the tumor, to invade the subventricular zones (SVZ) via the *corpus callosum* and to be resistant to radio- or chemo-therapy in a CXCR4/CXCL12-dependent manner^4,5^, properties that have been confirmed in human^6,7^. In addition, they maintain an immune-suppressive microenvironment which is beneficial for the tumor growth ^8^.

C-X-C chemokine receptor type 4 (CXCR4) and its ligand “Stromal cell-derived factor 1” (SDF1, also known as CXCL12) activate a signaling pathway emerging as a new marker of GSCs ^9^. CXCR4 is over expressed in many cancers including glioblastoma and its expression usually correlates with tumor progression ^10,11^. Most GSCs highly express CXCR4 and produce CXCL12, revealing an autocrine signaling loop ^12^.

Pre-clinical studies have shown that inhibition of CXCR4 is able to alter GBM characteristics of aggressiveness ^13,14^. Recently, a systemic delivery of nanoparticles coated with a CXCR4 antagonist (AMD3100 or Plerixafor), resulted in the inhibition of GBM growth and activation of an anti-tumor microenvironment, thus confirming the importance of the CXCL12/CXCR4 pathway as a therapeutic target ^13^. Several other CXCR4 antagonists have been evaluated in clinical trials for the GBM treatment, showing promising results when combined with standard therapies ^15^. Amongst the CXCR4/CXCL12 inhibitors, mutated forms of CXCL12 have been shown to efficiently antagonize CXCR4 signaling pathway. Crump *et al.* demonstrated that switching the CXCL12 second amino acid from a Proline to a Glycine (for that reason, called “P2G” in this manuscript), allows the mutated cytokine to bind CXCR4 while acting as a potent antagonist ^16^. The anti-cancer effect of P2G has been described in breast cancer and osteosarcoma murine models, with various impact on metastasis formation ^17,18^ but so far not in GBM.

Oncolytic Herpesviruses (oHSVs) represent one of the most promising anti-cancer tools in the evolving landscape of virotherapy. A deep knowledge of HSV genome and a relative flexibility for genetic modifications, have led to the engineering of a triple-mutated generation of oHSVs that has demonstrated its safety and is currently involved in clinical trials for the treatment of different tumors, including GBM ^19^. In 2021, the Japanese Pharmaceuticals and Medical Devices Agency (PMDA) conditionally approved the first oncolytic herpesvirus treatment (Delytact, G47Δ), for adult glioblastoma ^20^. However, in many cases, oHSVs are still not sufficient to counteract by themselves GBM malignancy. For this reason, they are either used as part of combination therapies, or armed to enable expression of a transgene that can amplify their anti-tumor activity, both approaches being a promising avenue.

We have engineered an oHSV armed with HA-P2G (called oHSV/P2G) which can replicate in the tumor and induce locally the production of P2G. This approach combines thereby the well-described effects of oHSV virotherapy with the inhibition of a crucial signaling pathway, while avoiding side effects of a systemic administration of a CXCR4-antagonist. The current study focuses on oHSV/P2G’s impact on both GSCs self-renewal and migration abilities, two important intrinsic GBM properties linked with GSCs functions. These features have been addressed both *in vitro* on human patient-derived GSCs and *in vivo* in a murine orthotopic xenograft GBM model ^4^.

## RESULTS

### Human GSCs express CXCR4 and CXCL12 at various levels

Five patient GSCs lines derived from primary (T08, T013, T018, GB138^21^) or recurrent GBM (T033) have been used in this study. Considering the heterogeneity of patient-derived GSCs, these cell lines have been characterized for their capacity to express both CXCR4 and its ligand CXCL12. Flow cytometry analysis revealed that apart T08 that are barely positive for CXCR4 expression, the other GSCs lines express CXCR4 at different levels **(Fig. S1A)**, with a very high expression in GB138 and T033. The amount of CXCL12 produced in the supernatant and measured by ELISA, showed that GB138 and T033 highly express CXCL12 while T08 secrete CXCL12 at a very low level **(Fig. S1B).** T013 and T018 express both CXCR4 and CXCL12 at an intermediate level. Taking into consideration their very low percentage of CXCR4^+^ cells and low CXCL12 expression, T08 were considered as negative control for the characterization of oHSV/P2G.

### oHSV/P2G can antagonize CXCR4 signaling

oHSV/P2G has been engineered as follows. P2G coding sequence, under the EF1α promoter, has been inserted in the ΔICP34.5/ΔICP6 oHSV backbone, just after the pICP6-eGFP coding sequence (kindly provided by Prof. EA Chiocca, Brigham and Women’s Hospital, Boston, MA, USA). Moreover, P2G gene has been flanked by the IL-2 signal peptide and HA-tag coding sequence, to promote its secretion and allow its detection respectively (**Fig. 1A**). To ensure its replication in GSCs, US12 gene coding for ICP47 which interferes with peptide presentation has been deleted in the oHSV backbone. This deletion puts ORF11, normally expressed as a Late protein under the control of the ORF47 promoter, allowing thereby its expression as an Immediate Early protein, partly restoring the infectivity of the oncolytic virus^22^.

**Figure 1:**
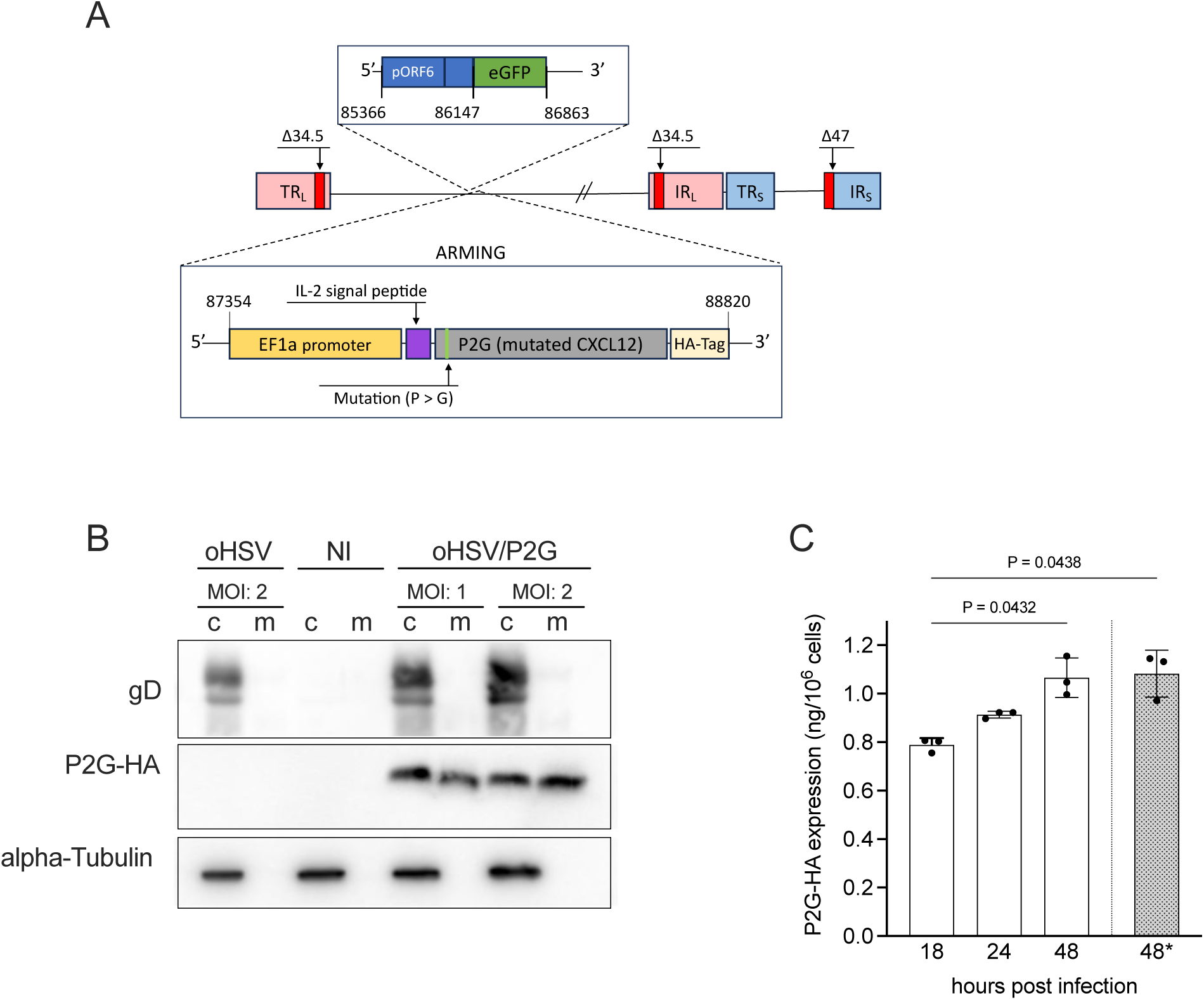
Construction and characterization of oHSV/P2G. **A.** Schematic representation of oHSV/P2G genome. **B.** P2G-HA expression by human GB138 primary cells infected by oHSV/P2G was analyzed by western blot analysis performed on non-infected (NI), oHSV- or oHSV/P2G-infected GB138 cells lysates (c) and supernatant (m). (Multiplicity of infection-MOI: 1 or 2). HSV glycoprotein D and α-tubulin detection were used as infection and loading control, respectively. **C.** P2G-HA secretion in the supernatant was quantified by ELISA. Human GB138 cells were infected with oHSV or oHSV/P2G (MOI : 1) and the supernatant was harvested 18, 24 or 48 hours post-infection (hpi). In addition, the supernatant harvested 48 hpi was filtered to eliminate viral particle (0.1 µm filtration indicated by *). Bars represent the mean (SD) of 3 independent experiments. Statistical significance was determined by one-way ANOVA, with Tukey’s multiple comparisons test with individual variance computed for each comparison.

Western blot (WB) analysis of oHSV- or oHSV/P2G-infected GB138 cells revealed that, as expected, oHSV/P2G infection led to the expression and secretion of HA-tagged P2G (**Fig.1B**). oHSV/P2G-infected T08 were able to efficiently express and secrete P2G as well (data not shown). Quantification of HA-tagged P2G in the supernatant of oHSV/P2G infected GB138 cultures confirmed that P2G-HA is increasingly secreted up to 48 hours after infection (**Fig.1C**). Filtration (0.1 µm filter) of the supernatant, previously proved effective in removing the viral particles without affecting the proteins in the medium ^23^, did not affect the amount of P2G-HA **(* in Fig. 1C)**, making it possible to evaluate P2G impact independently of the viral infection or virus-induced cell death. Importantly, the amount of P2G detected in the supernatant (± 1 ng/10^6^ GB138 cells) was in the same range as the amount of CXCL12, the endogenous ligand of CXCR4 (± 0,6 ng/10^6^ GB138 cells).

Insertion of P2G transgene into the fQuick backbone did not impair the capacity of the virus to replicate in human glioblastoma **(Fig. S2A)** and importantly, oHSV- or oHSV/P2G infection has a similar impact on cell proliferation **(Fig. S2B)**. Culture in the presence of oHSV- or oHSV/P2G-conditioned medium (cm) did not affect cell proliferation either (**Fig. S2C**). Finally, cell-death upon infection was similar with both viruses (**Fig. S2D**).

CXCR4 activation by CXCL12 triggers intracellular pathways, whose activation leads to the ERK and STAT3 phosphorylation ^24^. To confirm that P2G can antagonize CXCR4 signaling pathway, GB138 cells have been cultured in the presence of non-infected (NI), oHSV(cm) or oHSV/P2G(cm) and the levels of phosphorylation of ERK and STAT3 were evaluated by Western Blot **(Fig. 2)**. The densitometric quantification indicated that phosphorylation of both ERK and STAT3 was impaired in presence of oHSV/P2G(cm), whereas it was not modified in the presence of oHSV(cm). AMD3100, a well described CXCR4 inhibitor, showed a similar effect on ERK and STAT3 phosphorylation confirming the antagonistic properties of P2G.

**Figure 2:**
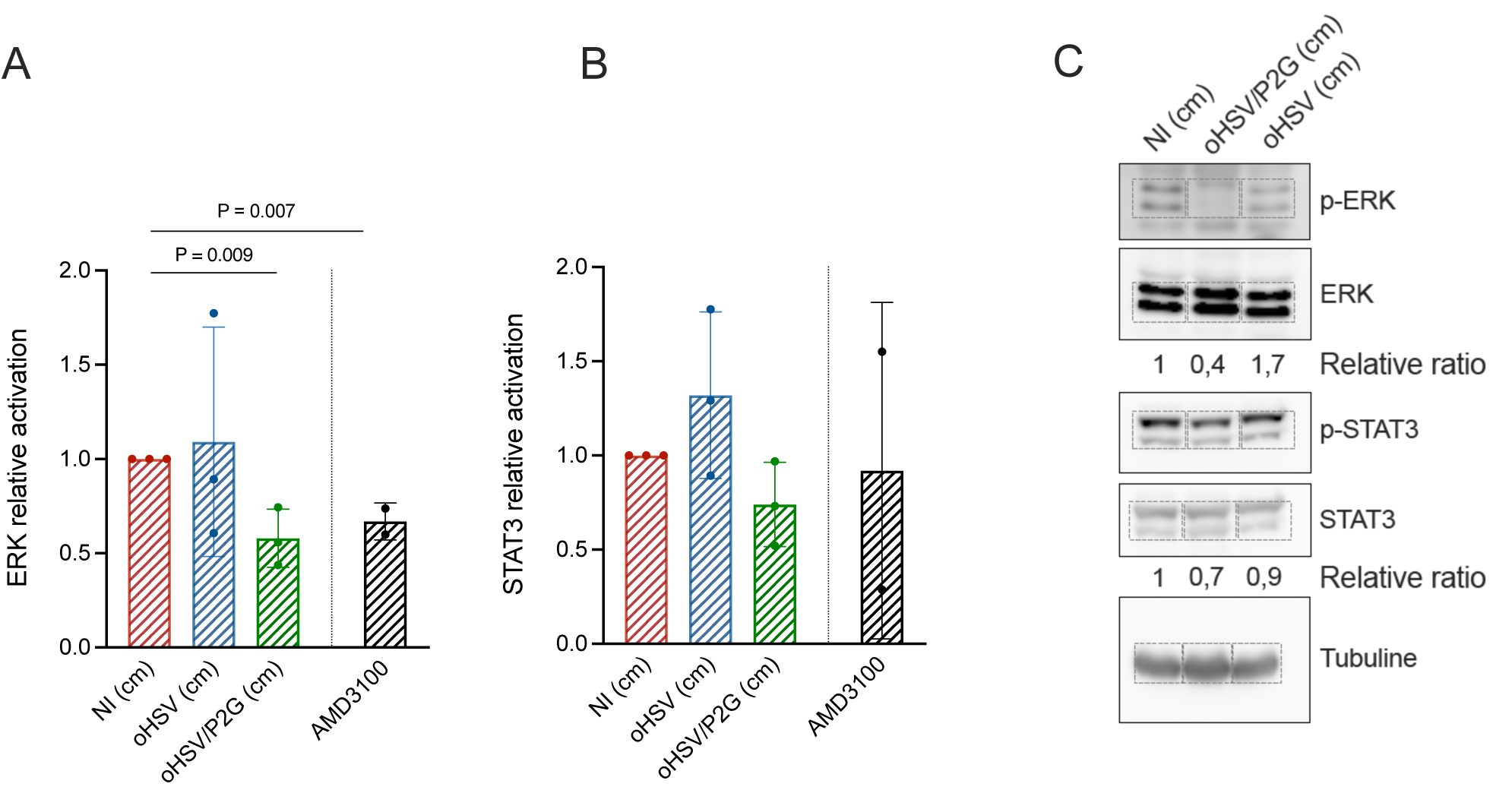
P2G impairs CXCR4 signaling pathway. P2G effect on CXCR4 signaling pathway was evaluated by western blot (WB) quantification of ERK **(A)** and STAT3 **(B)** phosphorylation in GB138 cells cultured with oHSV- or oHSV/P2G-conditioned media (cm). AMD31000 (Plerixafor) was used as a positive control of CXCR4 pathway inhibition. p-ERK/ERK and pSTAT3/STAT3 ratios were quantified by bands densitometry after western blotting and expressed as the relative ratio with the ratio in cells cultured with NI(cm) considered as 1. Bars represent the mean (SD) of 3 independent experiments. Statistical significance was determined by unpaired t test analyses. Representative picture of a WB is shown in **(C)**. Bands analyzed by densitometry for quantification are delineated by dotted lines.

### oHSV/P2G decreases GSCs stemness markers expression and counteracts GSCs self-renewal abilities

To investigate the effects of oHSV/P2G on GSCs key features, we first analyzed its impact on the expression of several neural stem cells markers, whose concomitant expression is generally linked to the presence and activity of GSCs ^25^. Although oHSV infection of GB138 increased INTα6 and CD44 or decreased SOX-2, OCT7 and SALL2 expression, oHSV/P2G infection led to a significant decrease of all these stemness markers expression except for SALL2 expression which decreased but not significantly **(Fig. 3A)**. Conversely, oHSV/P2G infection of T08 (CXCR4 ^low^) cells had no significant effect on the expression of any of the analyzed stemness markers (**Fig. 3B).** In line with these observations, infection of GB138 cells by oHSV/P2G decreased the number of CD133^+^ cells compared to NI cells as analyzed by flow cytometry and this effect was significantly more pronounced than upon oHSV-infection **(Fig. 3C).**

**Figure 3:**
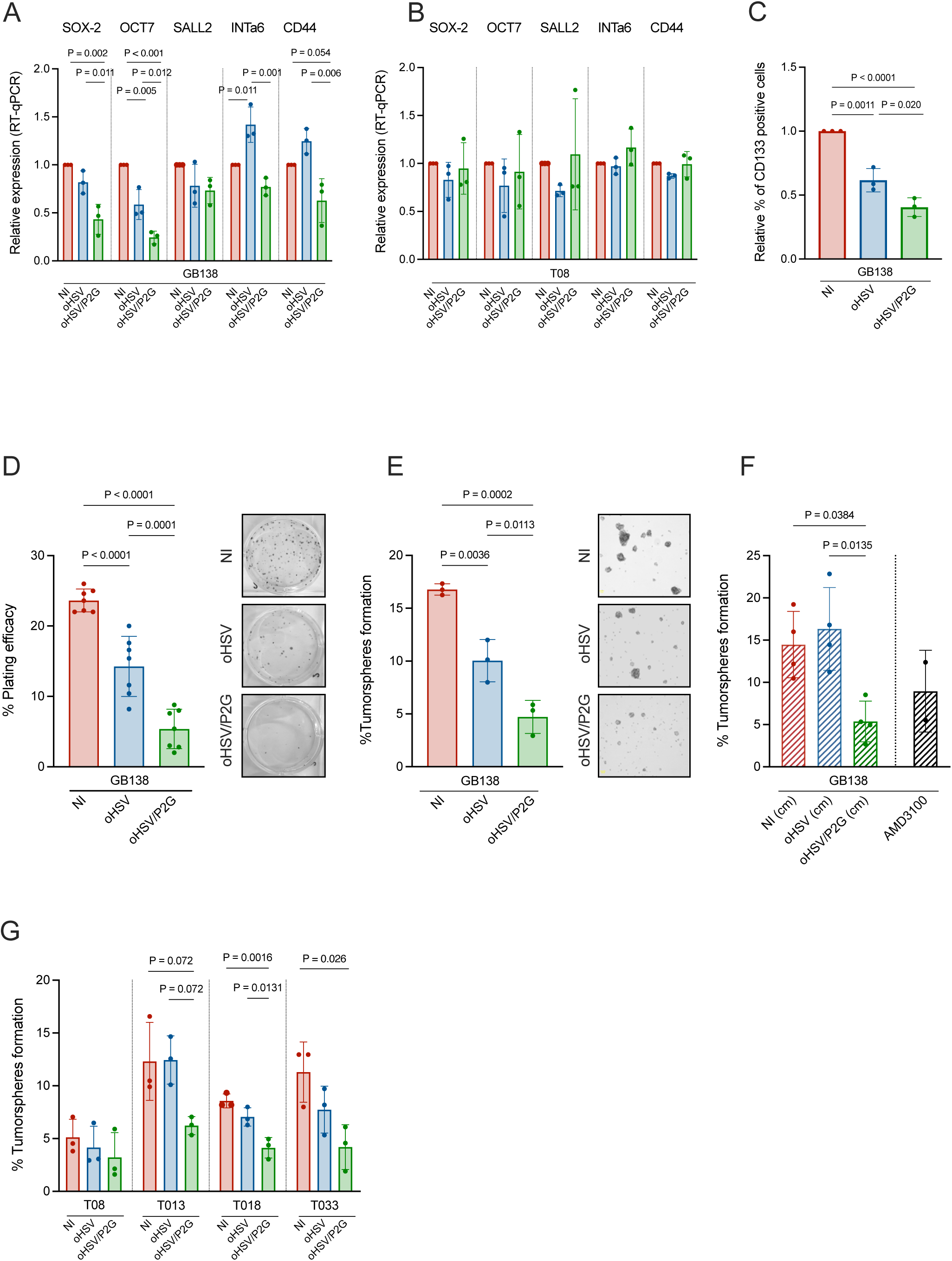
oHSV/P2G impacts stemness markers expression (A-C) and GSCs capacity to self-renew (D) and to form tumorspheres (E-G). **A-C:** Human GB138 **(A and C)** or T08 **(B)** primary cells were cultured as tumorspheres and infected with oHSV or oHSV/P2G (MOI 0.1). Expression of stemness markers (SOX2, OCT7, SALL2, INTα6 and CD44) was evaluated 6 days post-infection by RT-qPCR and expressed relative to the level of expression in non-infected cells considered as 1 **(A and B).** In parallel, GB138 cells were dissociated and the percentage of cells expressing CD133 at the plasma membrane was evaluated by flow cytometry **(C)**. Bars represent the mean (SD) of 3 independent analyses. Statistical significance was determined for each gene by ordinary one-way ANOVA with Tukey’s multiple comparisons test, with a single pooled variance. **D:** The capacity of GB138 cells to self-renew upon oHSV- or oHSV/P2G-infection (MOI = 0.1), was measured by a clonogenic assay. Bars represent the mean (SD) of 7 wells (3 independent experiments; 1 to 3 wells/condition in each experiment). Statistical significance was determined by one-way ANOVA with Tukey’s multiple comparisons test, with a single pooled variance. A representative picture of each experimental condition is shown in parallel. **E-G:** The capacity of GB138 cells to form tumorspheres upon oHSV- or oHSV/P2G-infection was expressed as a percentage of plated cells forming spheres 18hours after plating **(E)**. Representative pictures of tumorspheres upon mock-(NI), oHSV- or oHSV/P2G-infection are shown in parallel. The experiment was repeated with GB138 cells cultured in the presence of NI-, oHSV-, oHSV/P2G-conditioned media or NI media supplemented with AMD3100 (40 nM) **(F).** Finally, 4 other patient-derived primary GSCs (T08, T013, T018 and T033) were mock infected or infected with oHSV or oHSV/P2G **(G)**. Bars represent mean (SD) of 3 independent analyses. Statistical significance of E and F data was determined by one-way ANOVA with Tukey’s multiple comparison test with a single pooled variance.

The self-renewal ability of GB138 was then evaluated either by clonogenic or tumorsphere formation assays^26^, which respectively reflect the capacity of a single cell to form a colony by clonal expansion or to form spheres, two key features of GSCs. To limit the effect of the virus itself and avoid important cell death, cells were infected with a low MOI (MOI = 0.1). The clonogenic assay revealed that after 6 days of culture, the number of colonies was lower in the infected cells than in the control condition, indicating that the infection itself impairs the self-renewal capacity of GB138 which nevertheless was even significatively more affected upon oHSV/P2G infection (**Fig. 3D**).

To evaluate their capacity to form tumorsphere, GB138 cells were infected and cultured for 6 days in sphere-forming conditions. Again, the infection itself had an impact on the capacities of GB138 cells to form tumorspheres (**Fig. 3E**), but the number of tumorspheres observed in oHSV/P2G-infected cells was significantly lower than in oHSV-infected cells, demonstrating an effect of P2G itself. This assays, repeated in the presence of conditioned medium or AMD3100, confirmed that very few spheres were observed upon oHSV/P2G(cm) or AMD3100 treatment as compared to the number of spheres observed in NI or cells cultured with oHSV(cm) (**Fig. 3F**). These results were confirmed using the other GSCs, except for T08 for which a slight but not significant decrease was observed upon oHSV/P2G infection (**Fig. 3G**).

### oHSV/P2G counteracts GSCs migration *in vitro*

Another important GSCs feature observed *in vivo* is their capacity to escape the tumor mass and migrate to a stem cells niche by a CXCR4/CXCL12-dependent mechanism. This migration can be observed in the *corpus callosum,* through which they can reach the subventricular zone ^4,27^. This capacity to migrate to the surrounding tissues is one of the mechanisms underlying GBM recurrence. The capacity of oHSV/P2G to impair GSCs migration was first evaluated *in vitro* by transwell or sprouting assays. In these two assays, cell migration increased upon CXCL12 and decreased in presence of AMD3100, confirming that this feature is influenced by CXCR4/CXCL12 signaling (**Fig. S3**).

GB138 tumorspheres were infected for 18 hours with oHSV or oHSV/P2G (MOI 0.1) before being dissociated and seeded onto laminin-coated transwells. Counting of cells that had migrated throughout the membrane over a 48h period showed that the capacity of GB138 cells to invade the membrane was significantly impaired upon oHSV/P2G infection, compared to non-infected and even oHSV-infected cells **(Fig. 4A)**. In line with these results, the invasion ability of GB138 cells, cultured with oHSV/P2G(cm) or with AMD3100 was significantly lower than in presence of non-infected or oHSV(cm) **(Fig. 4B)**. These data were confirmed by crystal violet quantification (data not shown).

**Figure 4:**
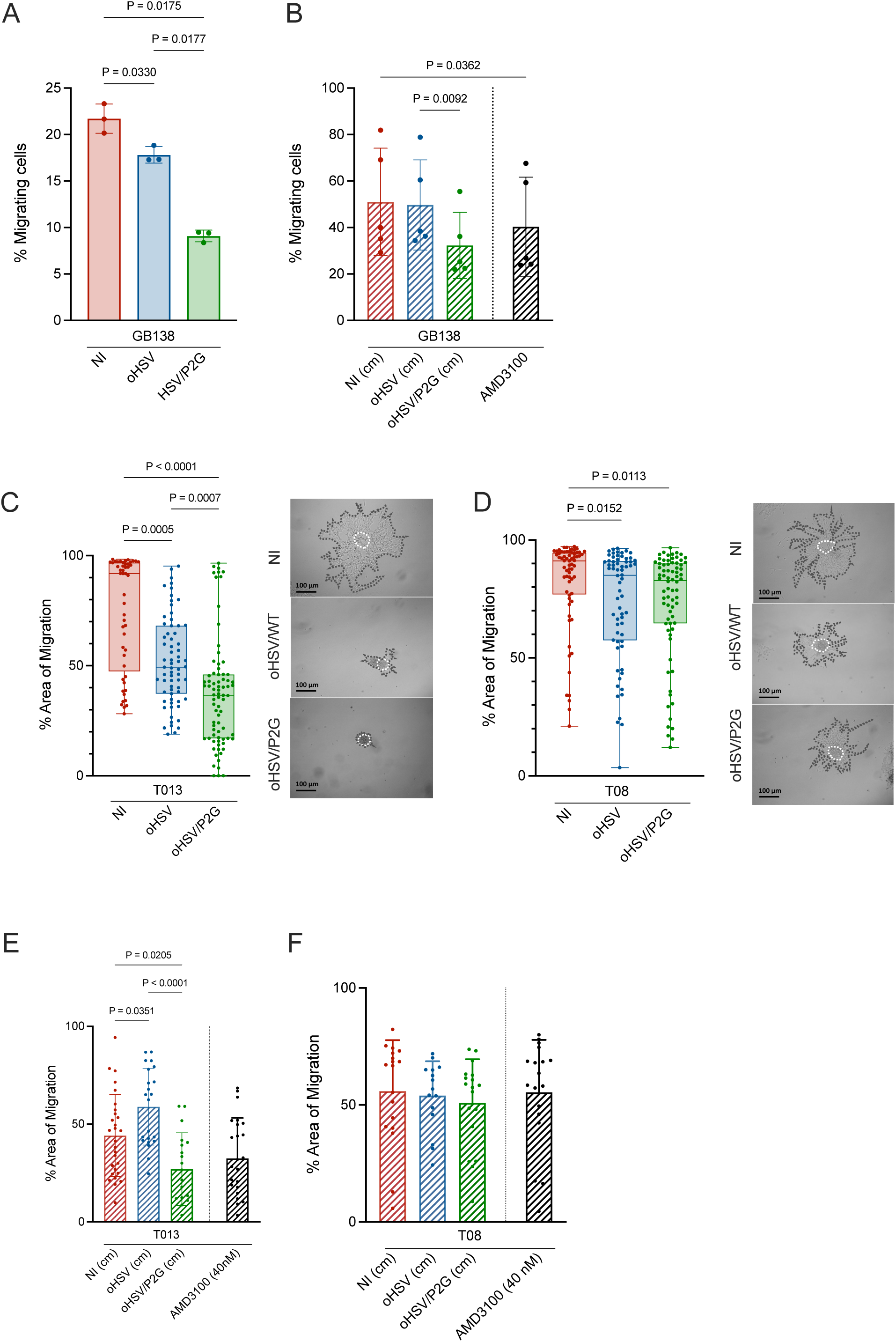
oHSV/P2G impacts GB cell migration. **A and B**: **Transwell assay**: GB138 cells infected or not with oHSV or oHSV/P2G (MOI: 0.1) were cultured in the upper insert of a two compartments culture device (transwell) and their capacity to migrate through the transwell membrane was evaluated after 48h **(A)**. The same experiment was performed with cells cultured in the presence of non-infected- (NI), oHSV- or oHSV/P2G-conditioned media (cm) or NI-conditioned media supplemented with AMD3100 (40 nM) **(B).** Bars represent the mean (SD) of 3 **(A)** or 5 **(B)** independent experiments. Statistical significance was determined by RM one-way ANOVA, with Tukey’s multiple comparisons test, with individual variance computed for each comparison. **C to F**: **Sprouting assay**: T013 (CXCR4^medium^) **(C)** and T08 (^Low^) **(D)** GSCs cultured as tumorspheres were either non infected or infected with oHSV or oHSV/P2G. Representative pictures are shown in parallel. The spheres areas were measured at 1 hpi (hours post-infection, white dotted lines) and 24 hpi (dark dotted line) and the percentage of migration was expressed as follows:

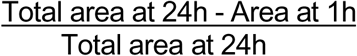 The same experiment was repeated with T013 **(E)** or T08 **(F)** cells cultured with NI-, oHSV- or oHSV/P2G-conditioned media or NI-conditioned media supplemented with AMD3100. Each dot represents one sphere while bars represent the mean (SD) of the spheres measured in each of the 4 (C and D) or 3 (E and F) independent experiments. Statistical significance was determined by Kruskal-Wallis test (C and D) or ordinary one-way ANOVA (E and F).

To confirm the oHSV/P2G effect on GSCs migration, sprouting assays were performed with T013 (CXCR4^medium^) and T08 (CXCR4^low^) tumorspheres previously infected or not with oHSV or oHSV/P2G. Twenty-four hours after seeding, migration of T013 cells was clearly impaired by oHSV/P2G infection while the effect on T08 was very limited **(Fig. 4C and D)**. Sprouting assay performed on cells cultured with conditioned media or media supplemented with AMD3100 **(Fig. 4E and F)** confirmed these observations. Although oHSV(cm) increased T013 migration, the presence of P2G or AMD3100 significantly decreased cell migration **(Fig. 4E)**. Conversely, migration of T08 cells was not affected neither by P2G or by AMD3100 **(Fig. 4F).**

### oHSV/P2G impairs tumor growth and GB cells migration *in vivo*

Finally, the capacity of oHSV/P2G to interfere with tumor development and migration was evaluated *in vivo* using an orthotopic xenograft murine model **(Fig. 5A).** The first experiment was performed with 9 mice (3 in each experimental group) and repeated with the same setting with 20 mice (6 for PBS or oHSV and 8 for oHSV/P2G treatments). Briefly, GB138-RFP^+^Luc^+^ cells were engrafted into the right striatum of *nude* mice (Day 0). On Day 19, *in vivo* bioluminescence imaging was performed to monitor tumor growth. Statistical differences between experimental groups were analyzed with R to take into consideration the potential bias resulting from two independent experiments **(R script in Supplemental material)** and showed that all groups harbor a tumor with comparable size, without any statistical difference **(Fig. 5B)**. On Day 20, PBS, oHSV or oHSV/P2G (1 x 10^6^ PFU) were injected into the tumor mass. Mice were sacrificed on Day 47 and brains were cleared for lightsheet microscopy analysis. The tumor volume estimation after 3D-reconstruction clearly indicated that tumors in the oHSV- and oHSV/P2G-treated mice were smaller than in the PBS group (**Fig. 5C)**. Importantly, while all PBS- or oHSV-treated mice harbor tumors, two oHSV/P2G-treated mice did not present any tumors at the end of the experiment.

**Figure 5:**
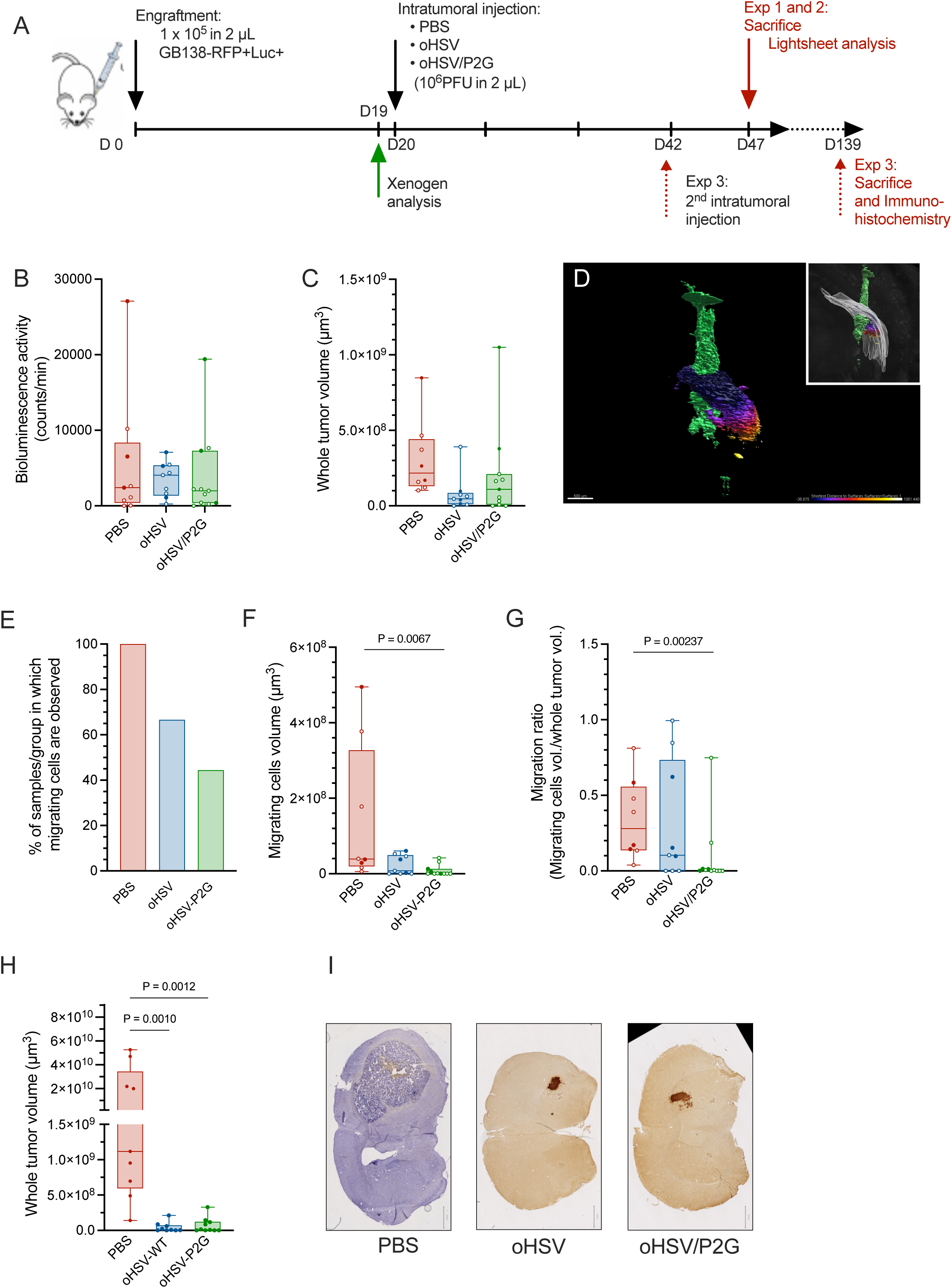
oHSV/P2G impairs GB cells migration in an orthotopic xenograft *in vivo* model. **(A)** Experimental settings of the orthotopic xenograft model. 1×10^5^ GB138-RFP^+^Luc^+^ cells were engrafted in Nude mice brain under stereotactic position (Day 0). One Day 19 (one day prior treatment), bioluminescence imaging analysis was performed to verify the presence and the size of the tumors. On Day 20, PBS, oHSV or oHSV/P2G (10^6^ PFU) were injected into the tumor mass under stereotactic control. (For the long-term assay, this treatment was repeated on Day 42). On Day 47 (Exp 1 and 2), brains were harvested and clarified for lightsheet imaging. For the long-term assay (Exp 3), mice were sacrificed on Day 139 for immunohistological analyses. Exp 1 (Plein dots) and 2 (Empty dots) results are illustrated in Figures B to G while Fig. H and I correspond to Exp 3 (Long term experiment). **(B)** Size of the tumor evaluated by bioluminescence imaging analysis on Day 19. **(C)** Whole tumor volume measured on tumor 3D-reconstruction (Imaris) of brains recovered on Day 47. **(D)** Representative picture of the 3D-reconstruction of the tumor mass (green) and of cells migrating (statistically fire-colored according to their distance to the central tumor mass) through the *corpus callosum* (grey, in the insert). Bar represents 500 µm. Pictures of Imaris 3D-modelization of all Exp 1 and 2 brains are shown in Figure S4. **(E)** Percentage of mice for which cells migrating through the corpus callosum were observed. **(F and G)** Volume of cells migrating towards the *corpus callosum* **(F)** or ratio of migrating cells regarding the volume of the whole tumor **(G**). Bars represent the mean (SD) of data from 8 (PBS), 9 (oHSV) and 11 (oHSV/P2G) mice. Statistical significance for Fig. B to G was determined in R according to the script available in Supplemental material. **(H and I)** Long term experiment (Exp 3). Whole tumor volume measured from immunohistochemical staining of serial sections of brains recovered on Day 139 **(H).** Bars represent mean (SD) form 9 (PBS and oHSV) and 10 (oHSV/P2G) mice. Statistical significance was determined by Kruskal-Wallis test. A presentative picture of one section is shown for each experimental group **(I)** Bar represents 1 mm.

In line with previous studies, lightsheet analyses confirmed that GBM cells mainly migrate from the tumor mass toward the *corpus callosum*. These cells were highlighted using fire statistical colors, reflecting migration distance relative to the tumor axis, and the volume of the infiltrative tumoral tissue was measured (Representative picture on **Fig. 5D**; all pictures in **Fig. S4**). While migrating cells were observed in all PBS-treated tumors, they were observed in 66% of the oHSV- and only in 44% of the oHSV/P2G-treated groups (**Fig. 5E**; **Fig. S4**). Moreover, the volume corresponding to cells migrating towards the *corpus callosum* was extremely low in treated mice and even significantly lower in oHSV/P2G-treated mice than in the control group (**Fig. 5F**). This was even more marked considering the ratio between the volume of migrating cells and the whole tumor volume (**Fig. 5G**).

To evaluate the capacity of oHSVs to limit tumor growth, a long-term experiment was performed. Mice were engrafted with GB138-RFP^+^Luc^+^ cells and treated with PBS (9 mice), oHSV (9 mice) or oHSV/P2G (10 mice) on Day 21 and on Day 42 (**Fig 5A**). On Day 139, no mice from any experimental group had died or even exhibited abnormal behavior warranting euthanasia. Mice were then sacrificed to evaluate tumor size (**Fig. 5H and I**). While most tumors of the PBS-treated mice were massive, some of them occupying almost the entire hemispheres, tumors in mice treated with oHSV or oHSV/P2G were either completely absent (3/9 and 4/10 in oHSV- and oHSV/P2G-treated mice respectively) or very small, demonstrating the virotherapy efficacy although in this immunocompromised model, the impact of oHSV and oHSV/P2G on tumor growth was similar.

## DISCUSSION

Glioblastoma is the most aggressive form of adult brain cancer and remains associated with poor prognosis mostly due to therapeutic failure and recurrences ^1^. Glioblastoma stem-like cells (GSCs) that express stemness markers, display strong self-renewal abilities and are generally resistant to most therapies, have been identified as contributing to GBM recurrences ^28^. Moreover, GSCs participate to the establishment of an immune-suppressive microenvironment sustaining tumor growth^8^. As shown in previous studies, some GSC features such as self-renewal, capacity to form tumorspheres or to migrate are mediated by the CXCR4/CXCL12 pathway ^9,12,29^. In many cancers including GBM, CXCR4 expression is correlated with poor prognosis, highlighting its inhibition as a potential therapeutic strategy^10,11^. Numerous preclinical studies have investigated the efficacy of systemic delivery of CXCR4 inhibitors with encouraging outcomes. However, systemic administration of CXCR4 inhibitors leads to side effects ^30^. Virotherapy and in particular herpesvirus virotherapy which has proven its safety for GBM treatment, constitutes a powerful tool to ensure a localized administration while limiting the side-effects.

We have engineered an oncolytic HSV-1 (Δγ34.5/ΔICP6/ΔICP47) armed with a mutated form of CXCL12 (called P2G) described in the literature as a potent CXCR4 inhibitor ^16^. The insertion of P2G does not impair virus replication and leads to the secretion of P2G by the infected cells. Furthermore, the amount of P2G secreted by the infected GBM cells is similar to the amount of endogenous CXCL12.

Although the virus itself influences human GSC phenotype and despite the fact that arming does not induce additional cell death compared to the virus itself, the presence of P2G has a significant impact on GSCs features. Arming indeed significantly decreases stemness markers expression, impairs self-renewal and migration capacities of GSCs as confirmed by culture with conditioned media devoid of viral particles. Importantly, P2G effect depends on the level of CXCR4 in patient-derived GSCs and is therefore very low in CXCR4^Low^ GSCs.

In an orthotopic xenograft murine model, intratumoral injection of either oHSV or oHSV/P2G significantly reduces tumor size, as estimated by 3D-reconstructed lighsheet microscopy images. It should be noted that in Exp 1, one of the oHSV/P2G-treated mice harbors a huge tumor, comparable to the tumors usually observed in the PBS group. This could account for technical issues, in particular to the difficulties to ensure the virus injection in the tumors which growth quite slowly and are very small at the time of treatment. Interestingly, while all PBS- or oHSV-treated mice developed a tumor, two (out of 11) oHSV/P2G-treated mice did not as observed in lightsheet microscopy at the end of the experiment (Day 47, Fig S4). Importantly, in a long-term experiment in which mice were treated twice (3 weeks apart) and sacrificed about 100 days after the second treatment, tumors in the PBS group were huge, occupying, in some cases, almost the entire hemisphere while tumors in both oHSV- and oHSV/P2G-treated mice were very small and even undetectable in 33% (3 out of 9) and 25% (4 out of 10) in oHSV- and oHSV/P2G-treated mice respectively. These observations demonstrate that virotherapy could lead to a long-lasting impact despite the absence of adaptive immune response in *nude* mice. Surprisingly, despite the size of the tumors, none of the PBS-treated mice lost weight or displayed abnormal behavior. However, for ethical reasons, all mice were sacrificed on Day 139, and no survival assay was performed. Notwithstanding encouraging results demonstrating the capacity of oHSV/P2G to significantly impair stemness features *in vitro*, we were not able to demonstrate *in vivo* a significant improvement upon oHSV/P2G treatment compared to non-armed oHSV. Considering that *in vitro,* oHSV and oHSV/P2G lead to cell death with the same efficacy and impact similarly cell proliferation, it is not surprising that *in vivo*, in absence of adaptive immune response known to play a critical role in the virotherapy efficacy, no significant differences of tumor size could be observed about 100 days after the viral injection. However, it is important to note that while most *in vitro* experiments were performed with a low MOI (0.1), a high dose of virus (10^6^ PFU) was injected in the tumor to ensure the virotherapy efficacy. Although impossible to evaluate, the MOI at the site of injection is thus probably much higher than in *in vitro* experiments. We could not rule out that, *in vivo*, the viral infection itself has such a strong impact on tumor growth that the impact of P2G is underestimated. It would be worth evaluating whether a lower MOI would have a less drastic effect on the tumor size while sustaining a stronger and long-lasting production of P2G that would highlight its impact.

Considering that GSCs escape the tumor through a CXCL12 gradient and migrate through the *corpus callosum* to the SVZ, on a CXCR4-dependant manner ^4,31^, leading to treatments failure and recurrences, impairing cell migration might thus be beneficial. We clearly showed, both *in vitro* and *in vivo,* that P2G-arming has a significant impact on cell migration. Importantly, *in vivo*, no tumor cell migration was observed in 5 mice (out of 9; 55%) in the oHSV/P2G group while observed in all PBS- and in 3 (out of 9; 33%) oHSV-treated mice. Moreover, considering the size of the tumors, cell migration was extremely limited in all oHSV/P2G samples. However, SVZ is not the only niche where GSCs can hide. Hira and colleagues have indeed shown that hypoxic peri-arteriolar tissue constitutes a GSC niche similar to the hypoxic peri-arteriolar hematopoietic stem cell niches ^32^ and that CXCR4 antagonists may mobilize GSCs from this niche and sensitize them to therapy^33^. Surgical resection and radiotherapy considered as first-line treatment, are usually followed by vasculogenesis promoted by a CXCR4-dependent recruitment of bone marrow derived cells (BDMCs) which can be inhibited by CXCR4 inhibitors. Although not investigated in the current study, it would really be worth investigating tumor vasculogenesis and mobilization of both GSCs and BMDC which would reinforce the rationale of oHSV/P2G administration after the current treatment.

Besides their capacity to lyse tumor cells, oncolytic viruses can also impact TME. CXCR4 inhibitors have been shown to reshape the tumor microenvironment (TME) and to potentiate immunotherapy in melanoma ^34^, in ovarian cancer ^35^ and in glioblastoma ^36^ ^37^. They can trigger the production of cyto- or chemokines and thereby recruit immune cells ^38–40^. In preclinical models, they have been shown to switch macrophages from a pro-tumor M2-like to an anti-tumor M1-like phenotype ^41^. Of note, pro-tumoral macrophages, particularly abundant in GBM, correlate with faster progression and therapy resistance. On the other hand, CXCR4 inhibition has recently been shown to decrease Treg recruitment, to impair their functions and more generally to counteract immunosuppressive properties of the TME ^42^. It is thus crucial to evaluate the oHSV/P2G effect on the immune response. Characterization of oHSV/P2G in an immunocompetent syngeneic model will allow to confirm that in GBM, CXCR4/CXCL12 inhibition not only modify the GSCs phenotype, impairs their capacity to migrate but also modulates the TME participating thereby to a better tumor control as described for other cancers.

Altogether, our results demonstrate that local expression of P2G through virotherapy has a significant impact on GSC phenotype, reducing their stemness markers expression and significantly impairing their capacity to self-renew and to migrate both *in vitro* and *in vivo*. oHSV/P2G-virotherapy might thus be considered as an add-on to the first-line treatment, to limit the risk of relapse and improve GBM prognosis.

## MATERIAL and METHODS

### Cell lines

VERO cells (ATCC, #CCL-81) and patient-derived human glioblastoma GB138 cells ^21^, were maintained in Dulbecco’s modified Eagle minimal essential medium high glucose (DMEM HG, Lonza, Verviers, Belgium) supplemented with 10% fetal bovine serum (FBS). GBM patient-derived cultures (T08, T013, T018 and T033^21^) were established from freshly resected glioblastoma tissue obtained from GBM patients. They were cultured as tumorspheres in stem cell medium (DMEM/F-12 with GlutaMAX (Gibco) supplemented with B27 (1/50) without vitamin A (Gibco), 1% Penicillin-streptomycin (Lonza, Verviers, Belgium), 1 μg/ml of heparin (n 7692.1, Carl Roth, Belgium), human EGF (20 ng/ml) and βFGF (20 ng/ml) (Peprotech). When indicated, GB138 were cultured as monolayers.

### Construction of recombinant oHSVs

Recombinant viruses were engineered in fHsvQuik-1 Bacterial artificial chromosome (BAC) containing an attenuated HSV-1 (strain F) (Δγ34.5, ΔUL39, GFP^+^; kind gift from A. Chiocca, Brigham and Women’s Hospital, Boston, MA, USA). Further ICP47 deletion was done as described by Todo T *et al*., 2001 ^43^. Recombinants were obtained by the two-step Red recombination technique “en passant” ^44^. The sequence of human mutated CXCL12 sequence (second amino acid mutated P to G) flanked by the IL-2 signal peptide and HA-Tag in 5’ and 3’ respectively and controlled by the EF1a promoter was inserted just after the eGFP gene (**Fig. 1A**). Vero cells seeded in 6-well plates were transfected with this construct using JETPEI (Polyplus, Illkirch – FRANCE). Virus stocks were produced as previously described ^45^. Viral particles were then ultracentrifuged, resuspended in PBS with 10% glycerol and stored at -80°C. Virus titration was performed in VERO by plaque assay^46^.

### Virus and conditioned media production

GB138 cells were grown in DMEM HG 10% FBS. At confluence, the culture medium was removed and replaced by DMEM/F12 (without added growth factors), and cells were mock-infected or infected with oHSV or oHSV/P2G (MOI: 1). After 48h, media were collected, centrifuged (260g, 5 min), filtered with 0,1 µm filter (Pall Life Sciences® 4611 Acrodisc® Syringe Filters with Supor® Membrane, 25mm, 0.1um, Sterile), aliquoted and stored at -80°C.

### Viral growth assay

50k GB138 cells were seeded in 24-well plate for 24h and then infected with oHSV or oHSV/P2G (MOI 0.1 and 1). Virus replication was measured by recording GFP signal for 48h (Incucyte® S3). Total green object area (µm^2^/well) was used to assess virus replication. Total object area measured with contrast imaging was recorded in parallel and expressed relative at T0 to evaluate the cell proliferation.

### Detection of HA/P2G by Western blot

400K cells were seeded for 24h before being infected with oHSV or oHSV/P2G (MOI 1 or 2). Supernatant and cells were harvested 18hours post infection (hpi).

Cells were lysed using modified RIPA buffer (Tris-HCl 50mM pH7.5, NaCl 150mM, EDTA 1mM, NP40 1%, DOC 0,25%) supplemented with proteases inhibitors (Complete, Roche). Supernatants were diluted in acetone (1:4) to precipitate the proteins and further centrifuged at 13 000 RPM for 30 min.

After electrophoresis and transfer onto PVDF membrane (GE Healthcare), membranes were incubated overnight at 4°C with anti-gD (Ref. sc-21719, Santa Cruz; dilution 1:1000), and anti-HA (Ref. ab9110, Abcam; 1:1000) antibodies. Mouse anti-alpha-tubulin (Ref. T6199, Sigma; 1:2000) was used as loading control. HRP-conjugated anti-mouse IgG (Ref. 7076, Cell Signaling; dilution 1:2500) or HRP-conjugated-anti-rabbit IgG (Ref. 7074, Cell Signaling; 1:2500) were used as secondary antibodies. After incubation with ECL (luminol: 25 mg/100 ml in Tris-HCl pH 8.6 0.1M; para-hydroxy coumaric acid 11 mg/10 ml DMSO; H202 35%), chemiluminescence signals were recorded with LAS4000 CCD camera (GE Healthcare).

### p-ERK And p-STAT relative quantification

125k GB138 cells were seeded in T25 in stem cell culture medium. After 6 days, the culture medium was replaced by conditioned media supplemented with CXCL12 (80 pM) to induce CXCR4 signaling pathway. After 30 min, cells were harvested, lysed using modified RIPA buffer and analyzed by western-blotting with anti-ERK (#9102, Cell signaling; dilution 1:2000), anti-pERK (#9106S, Cell signaling, 1:2000), anti-STAT3 (#9139 Cell signaling, 1:1000) or anti-pSTAT3 (#9145, Cell signaling, 1:1000) antibodies. Mouse anti-alpha-tubulin (#T6199, Sigma; 1:2000) was used as loading control. HRP signals were revealed using ECL and imaged with LAS4000 CCD camera (GE Healthcare). Bands densitometry was analyzed using ImageJ.

### HA-P2G quantification by ELISA

HA-P2G quantification in the culture supernatants was performed by ELISA with the HA tag ELISA kit (Ref MBS3802035, MyBioSource), according to the manufacturer protocol.

### Resazurin assay

50k GB138 cells were seeded in 24-well plate for 24h and then infected with oHSV or oHSV/P2G (MOI 1). After 24, 48 or 72h, culture medium was removed, cells were washed with PBS, and medium were replaced by 300 µL of resazurin solution (1/5 resazurin + 4/5 DMEM/F12). After 4h, resazurin was transferred into a 96-well black opaque plate. Fluorescence was measured using FilterMax F5 Multi-Mode Microplate Reader and expressed as relative fluorescent units (RFU, Ex = 530–570 nm, Em = 590–620 nm).

### RT-qPCR

Cells were seeded in 6-well plates (400K cells/well) for 24h before being infected with oHSV or oHSV/P2G (MOI: 0.1) for 18 hours and kept growing as tumorspheres.

Tumorospheres were then dissociated with Accutase (Biowest, Nuaillé, France), and 125K cells were seeded in T25 in stem cells medium. Six days after plating, the spheres were collected and total RNA was purified (Nucleospin® kit, Macherey-Nagel, according to the manufacturer’s protocol). One µg of RNA was reverse transcribed using RevertAid H Minus First Strand cDNA Synthesis Kit (Thermo Scientific) with Random primers. TBP or 18S were used as housekeeping genes. qPCR reaction samples were prepared as follows: 4 µl of the diluted cDNA (10 ng in total) were mixed with 5 µl of SYBR green (TAKYON, Eurogentec, Liege, Belgium) and 0.5 µL of each primer (4 µM each) in a final volume of 10 µl. Primers used for transcripts detection are described in Table 1. Quantitative real-time PCR was performed using the Roche LightCycler480® (3 min. at 95°C of activation; 45 cycles: Denaturation 95°C, 3 sec, Hybridization and Elongation 60°C 25 sec).

**Table 1:**
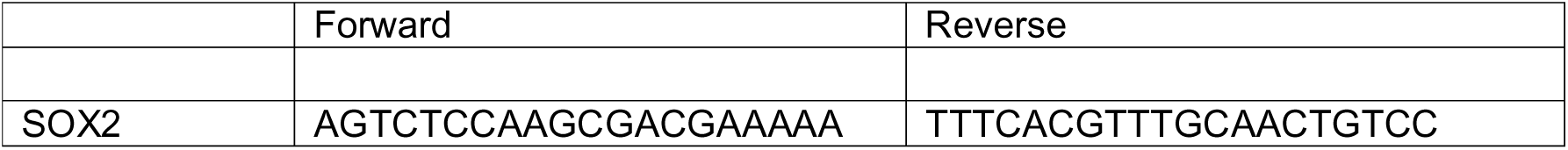
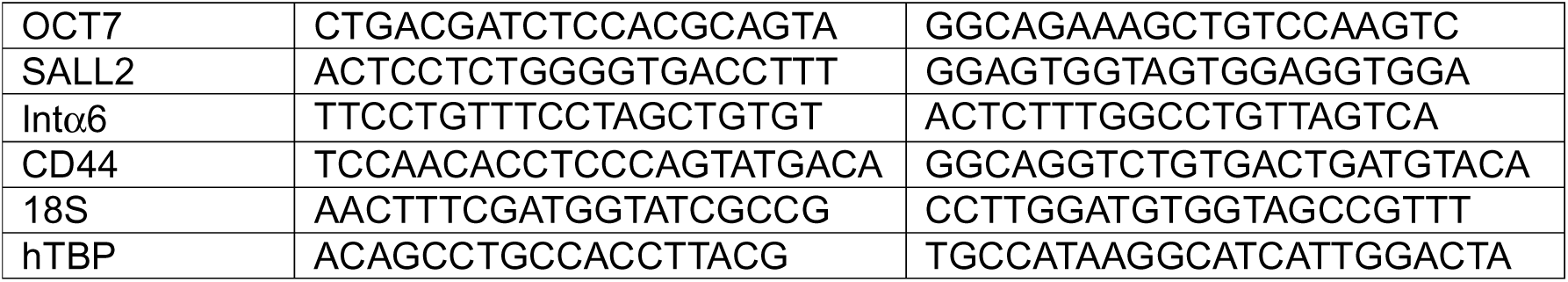
Primers used for RT-qPCR

### Flow cytometry

Tumorspheres or cells cultured as monolayers were washed with PBS and dissociated by incubation during 10 min at 37°C with Accutase or trypsin-EDTA (Biowest), respectively, then centrifuged (350g, 5 min, 4°C) and washed with Flow Buffer. 5 µl of APC-conjugated anti-CXCR4 (1/20; Biolegend, Amsterdam, The Nertherlands) or BV421-conjugated anti-CD133 antibody (1/25; BioLegend, Amsterdam, The Nertherlands) were added to 1×10^5^ cells in 100µl of Flow buffer and kept at 4°C for 1 hour in the dark. Cells were then washed and centrifuged at 400g for 4 min at 4°C. After a second wash, cells were resuspended in 200 µl of Flow buffer and analyzed with the FACS CANTO II (BD biosciences). Data were analyzed with FlowJo software.

### Clonogenic assay

GB138 cells were seeded in 6 wells-plates (400K cells/well) for 24 hours. Cells were then mock-infected or infected with oHSV or oHSV/P2G (MOI: 0.1) for 18 hours before being harvested and counted. 500 cells were re-seeded in 6-wells plates for 7 days. They were then fixed with paraformaldehyde (4%, 10 min, RT) and stained with Crystal violet (10 min, RT). After a final wash, colonies were counted, and the plating efficiency expressed as the number of colonies/ the number of plated cells *100. Finally, cells were lysed with 0.1% SDS and crystal violet absorbance was measured to confirm the colonies count.

### Tumorspheres assay

GB138 cells were seeded in 6 wells-plates in DMEM HG media + 10% FBS (400,000 cells/well) for 24h before being infected (MOI of 0.1). T08, T013, T018 and T033 cells were seeded in T25 in stem cell media (250k cells/flask). As spheres reach sufficient size, they were infected (MOI of 0.1). After 18h of infection, cells were either harvested with trypsin-EDTA or dissociated using Accutase. 125k cells were transferred to a T25 in stem cell media. Alternatively, non-infected cells were seeded (125k cells in a T25) in stem cell media in the presence of conditioned media. Tumorsphere formation efficiency was evaluated after 6 days and expressed as the number of tumorspheres/the number of plated cells *100.

### Migration quantification by Transwell assay

The upper chamber of ThinCerts™ transwells (pore size: 8µm; Greiner bio-one, cellstar®) placed in a 24-well plate (Greiner bio-one) were filed with 60 μL of laminin (20 µg/mL in PBS) and incubated for 24h at 37°C. Excess laminin was removed and 25k cells were plated in a final volume of 200 µl of conditioned media without growth factors and 300 µl of complete medium were added. After 48h, the medium was removed, and inserts were washed with PBS. Cells were then fixed with 4% paraformaldehyde for 10 min and washed with PBS, before being stained with crystal violet for 10 min. Non-migrating cells were removed from the transwell with a cotton swab and invading cells were counted with Image J on five pictures taken randomly (Magnification X20). The percentage of migrating cells is expressed as the number of invading cells/ the number of plated cells*100.

### Sprouting assay

Tumorspheres were either infected with oHSV or oHSV/P2G or cultured with conditioned media for 24h. They were then seeded into wells of a 96-well plate previously coated with 50 μL hydrobromide poly-D-lysine (10 μg/mL) for 30 min, washed with sterile water and left to dry overnight under the hood. If infected, tumorspheres were kept in DMEM media without growth factors while if pre-treated with conditioned media, they were kept in DMEM supplemented with conditioned media (50/50).

Pictures of the tumorspheres were taken with an optical microscope one hour after plating and after 24 hours of incubation. The migration level is measured using ImageJ and expressed as the percentage of migration regarding the tumorspheres area one hour after plating (Area at 24ℎ− area 1ℎ/Area 24ℎ∗100).

### *In vivo* experiments

All *in vivo* experiments were performed as previously approved by the Animal Ethical Committee of the University of Liège, in accordance with the Declaration of Helsinki and following the guidelines of the Belgium Ministry of Agriculture in agreement with European Commission Laboratory Animal Care and Use Regulation. Six weeks old female immunodeficient Crl:NU-Foxn1nu mice (Charles River Laboratories, Brussels, Belgium) were used for xenograft experiments. They were housed at the Animal Facility, University of Liège, in sterilized, filter-topped cages (controlled temperature: 22°C, controlled lighting: 12h day/night). After one week acclimatization period, intra-striatal grafts were performed following the previously described procedures^21^. Briefly, 100K GB138-RFP^+^Luc^+^ cells resuspended in 2 µl of PBS were injected into the right striatum (stereotactic coordinates: 0.5 mm anterior and 2 mm lateral from the bregma and at a depth of 2.5 mm) of mice previously anesthetized with an intraperitoneal injection (i.p.) of a Rompun (Sedativum 2%, Bayer, Brussels, Belgium) and Ketalar (Ketamin 50 mg/mL, Pfizer, Brussels, Belgium) solution (V/V) prepared just before injection. On Day 20 (all experiments) and on Day 42 (Exp 3), PBS (control) or oncolytic viruses (oHSV or oHSV/P2G; 10^6^ PFU in 2µl of PBS) were injected within the tumor, under anesthesia and using the same stereotactic coordinates. Mice were sacrificed on Day 47 (Exp 1 and 2; Lightsheet microscopy) or on Day 139 (Exp 3, Long-term experiment). Mice health status (weight and behavior) was evaluated daily.

ARRIVE 2.0 reporting guideline was used to assure the adequate management of animals^47^.

### Bioluminescence activity

Mice engrafted with GB138-RFP^+^Luc^+^ were injected i.p. with Beetle Luciferin Potassium salt (Ref. E1605, Promega) (150 mg/kg) and imaged (under 2.5% isoflurane anesthesia) with camera-based bioluminescence imaging system (Xenogen IVIS 50^®^; exposure time 1 min, 15 min after intraperitoneal injection). Regions of interest were defined manually, and images were processed using Living Image and Igor Pro Software (Version 2.60.1). Raw data were expressed as total counts/min.

### Brain tissue clarification and Light sheet microscopy acquisition and analysis

Mice were euthanized with i.p. injection of Euthasol Vet (140 mg/kg), followed by an intracardiac perfusion of ice-cold saline solution. Brains were further clarified to remove lipids and replace them with polyacrylamide. Afterwards, brains were incubated for 6h at RT in 50% RIMS solution (#D2158, Sigma-Aldrich; Refraction index: 1.46; 50% in water). 50% RIMS solution was then discarded and replaced by 100% RIMS solution for an overnight incubation at RT.

Eventually, brains were imaged using lightsheet microscopy. Images were acquired with 5X objective (zoom = 0.6; pixel size = 1.52; lightsheet 5.826 µm; center thickness = 12.4; image size = 2922.4×2922.4 µm^2^) and processed to obtain one merged image per plane and 3D-reconstruction using Zeiss Arivis software.

Tumors were visualized thanks to the RFP signal while the *corpus callosum* was annotated on each individual image based on the brain autofluorescence signal. Tumor and *corpus callosum* 3D-modeling were performed on Imaris Image Analysis Software and allowed whole tumor volume and migrating cells volume measurements.

### Statistical analysis

All statistical analyses were performed using GraphPad Prism 10. Data are displayed as Mean ± SD. Normality was evaluated using Shapiro-Wild test. Depending on the experiments, paired t-Test, one-way ANOVAs were performed as indicated in the figure legends. For in vivo experiments 1 and 2 (Fig 5), statistical significance was analyzed with R to take into consideration the putative experimental bias. R script is available in supplemental material.

## DATA AVAILABILITY SATEMENT

All original data shown in the figures and supplemental figures are available upon reasonable request

## Supporting information

Supplemental material: R script

## ACKNOWLEDGMENTS

PD and JSG have benefited respectively from a Postdoctoral and doctoral fellowship from TELEVIE-FNRS, Belgium. MD is a Research Fellow of the FNRS-Belgium. This work was supported by grants from the National Fund for Scientific Research (FNRS, Télévie); the Special Funds of the University of Liège; the Leon Fredericq Foundation, Liège,Belgium. The authors would like to thank Prof. A. Chiocca (Brigham and Women’s Hospital, Boston, MA, USA) for fHsvQuik-1 BAC, C. Desmet (GIGA, University of Liège) for his help for R-statistical analysis and all the members of the GIGA Viral Vector, Imaging and Flow Cytometry, Genomics platforms, and animal facilities for valuable technical support. We warmly thank Adeline Deward (www.illuminesciences.be) for the graphical abstract.

## AUTHORS CONTRIBUTIONS

Conceptualization: P.D., M.D. and C.S-D. Virus engineering: J.S-G, M.D. and P.D. with the help of M.L. *In vitro* experiments: P.D., M.D. *In vivo* experiments: M.D. with the help of A.L., C.L. and B.B. Manuscript writing: P.D., M.D., and C.S-D. Scientific discussions during the project: P.D., M.D., B.R., V.N., A.L., M.L. and C.S-D. All authors critically reviewed and edited the manuscript. Funding acquisition: C.S-D.

P.D. and M.D. contributed equally as first authors.

M.L. and C.S-D contributed equally as last authors.

All authors approved the final manuscript.

## DECLARATION OF INTERESTS STATEMENT

J.S-G is currently a Postdoctoral Research Fellow at the Brain Tumor Research Center, Massachusetts General Hospital (MGH), Boston, MA, USA

P.D. is currently scientist at Janssen Vaccines and Prevention B.V., Leiden, The Netherlands

The other authors declare no competing interests

