## Supplemental Figures for "An oncolytic herpesvirus expressing a CXCR4 antagonist interferes with Glioblastoma cells stemness features and migration"

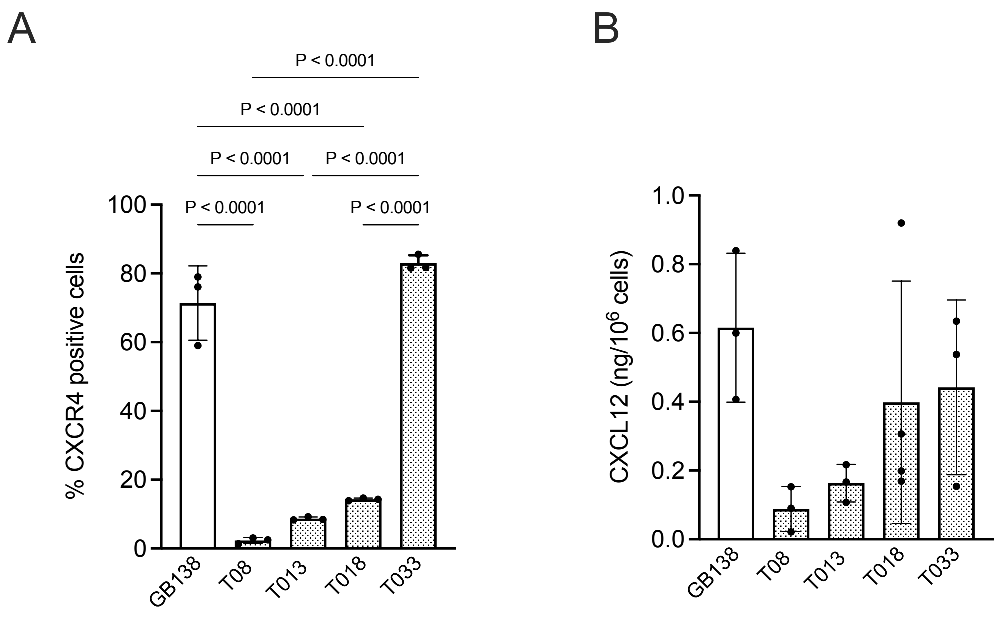


**Figure S1. CXCR4 and CXCL12 expression by patient-derived GSCs.**

CXCR4 **(A)** and CXCL12 **(B)** expression were evaluated on patient-derived GSCs (GB138, T08, T013, T018 and T033) cultured for 7 days either as monolayers (GB138) or as tumorspheres (T08, 13, 18 and 33). For CXCR4 analysis, cells were dissociated and the percentage of $CXCR4^{+}$ cells was evaluated by flow cytometry **(A).** In parallel, the cell culture supernatant was harvested for CXCL12 quantification by ELISA **(B)**.

Bars represent the mean (SD) of 3 independent experiments. Statistical significance was determined by ordinary one-way ANOVA with Tukey's multiple comparisons test, with a single pooled variance.

Figure S2

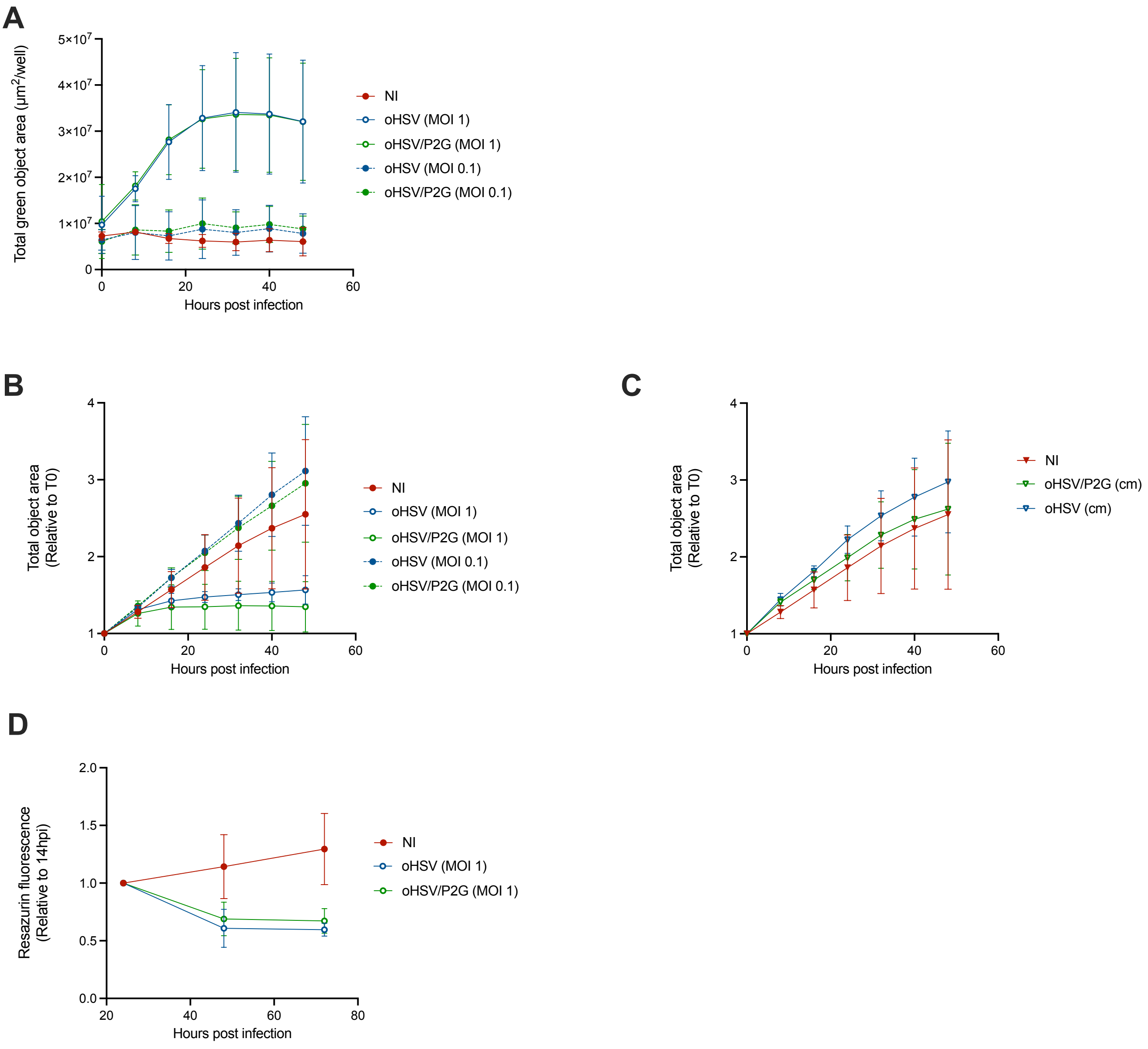


**Figure S2. P2G-HA expression does not influence viral replication, cells proliferation or cell-death**

Human GB138 primary cells were infected with oHSV or oHSV/P2G at MOI 0.1 or 1.

**(A)** The capacity of both viruses to replicate in GB138 cells was followed over a 48h period using Incucyte® S3 Live-Cell Analysis System recording eGFP expression. Total green object area (µm$^2$/well) was used to account for the spread of the infection.

**(B and C)** The capacity of GB138 cells to proliferate upon oHSV or oHSV/P2G infection **(B)** or cultured in presence of conditioned media **(C)** was followed over a 48h period using Incucyte® S3 Live-Cell Analysis System. Total object area (µm$^2$/well) was measured on phase contrast pictures and expressed relative to T0 considered as 1.

**(D)** GB138 Cell-death upon oHSV or oHSV/P2G infection was measured over a 72h period by resazurine assay and expressed as relative to fluorescence at 24hpi considered as 1.

Bars represent the mean (SD) of 3 independent experiments. Statistical significance was determined by Two-way ANOVA. No statistical differences between oHSV and oHSV/P2G were observed in any of the assays.

Figure S3

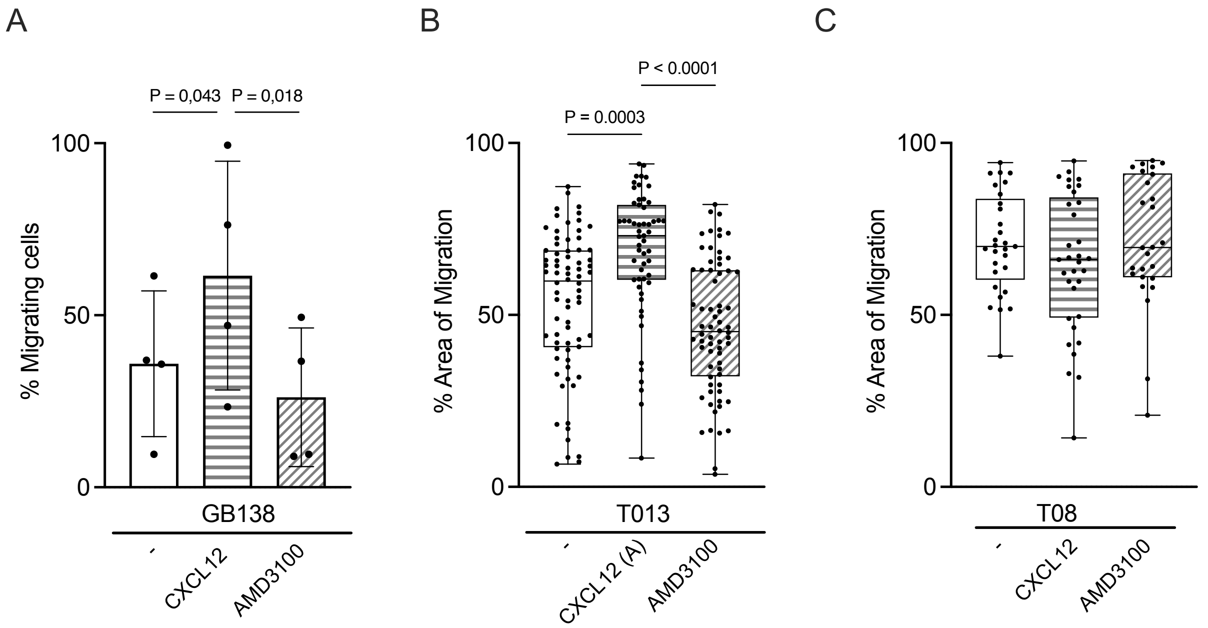


**Figure S3: GSC migration is a CXCR-dependent process.**

**(A):** Dissociated GB138 cells were cultured in two-compartments chambers in normal media or in media supplemented with purified CXCL12 (20 pM) or AMD3100 (40 nM).

The number of cells migrating through the transwell membrane was quantified by visual counting. Bars represents the mean (SD) of 4 independent experiments.

**(B and C):** T013 ($CXCR4^{Medium}$) or T08 ($CXCR4^{Low}$) tumorspheres were cultured in normal media or in media supplemented with CXCL12 (20 pM) or AMD3100 (40 nM).

The spheres areas were measured at 1 hpi and 24 hpi and the percentage of migration was expressed as follows:

$$\frac{\text{Total area at 24h - Area at 1h}}{\text{Total area at 24h}}$$

Each dot represents one sphere while bars represent the mean (SD) of all the spheres measured in 5 (B) or 3 (F) independent experiments. Statistical significance was determined by Kruskal-Wallis test.

Figure S4-A

*In vivo* experiment #1

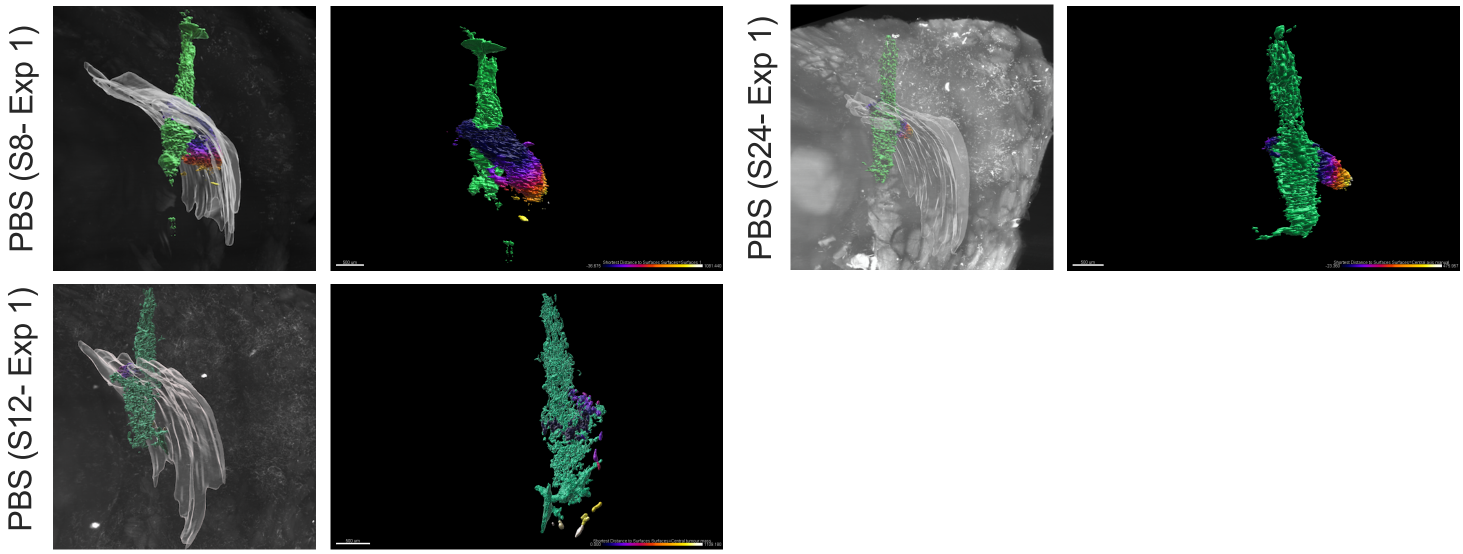


*In vivo* experiment #2

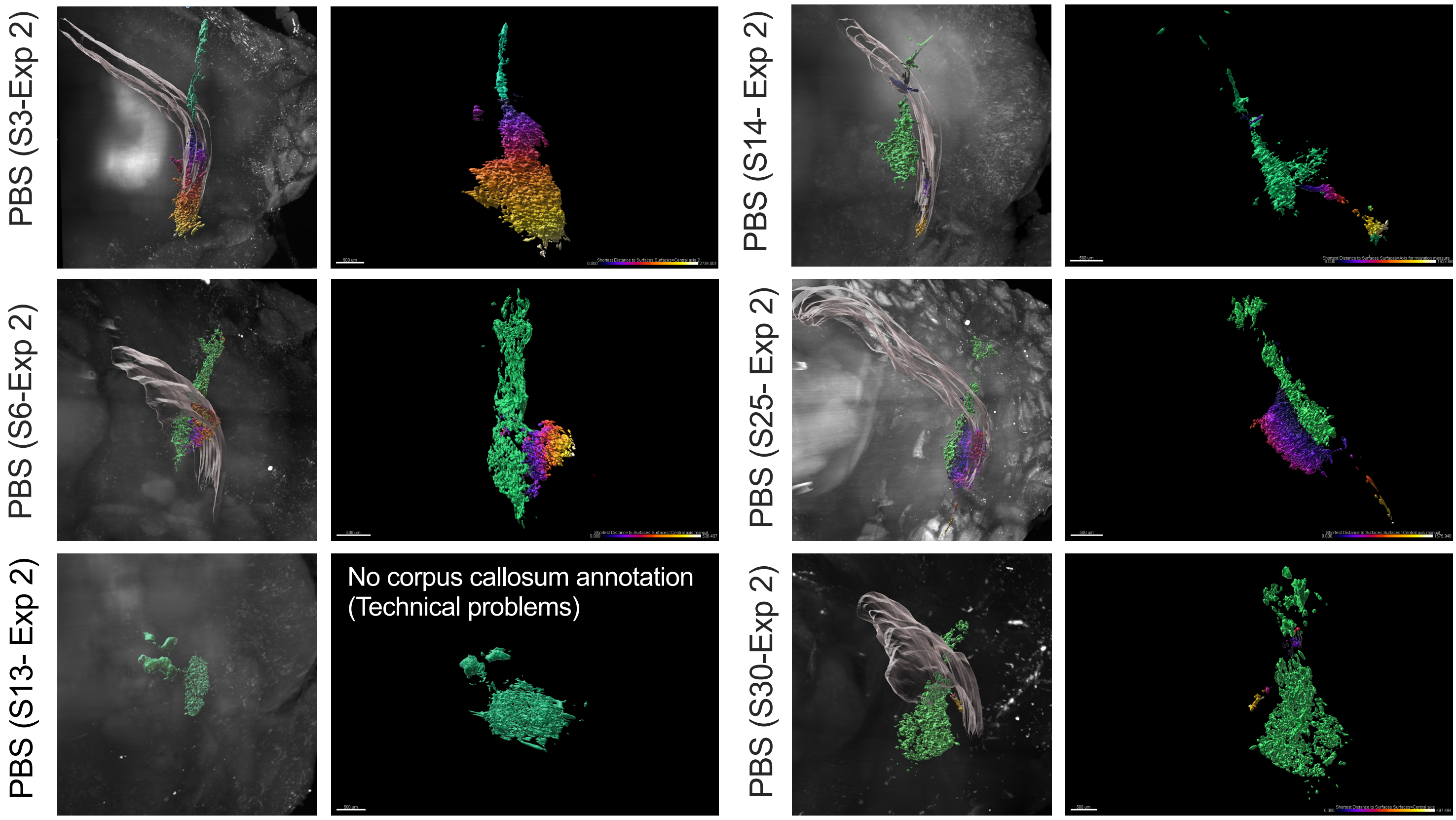

Figure S4-B

*In vivo* experiment #1

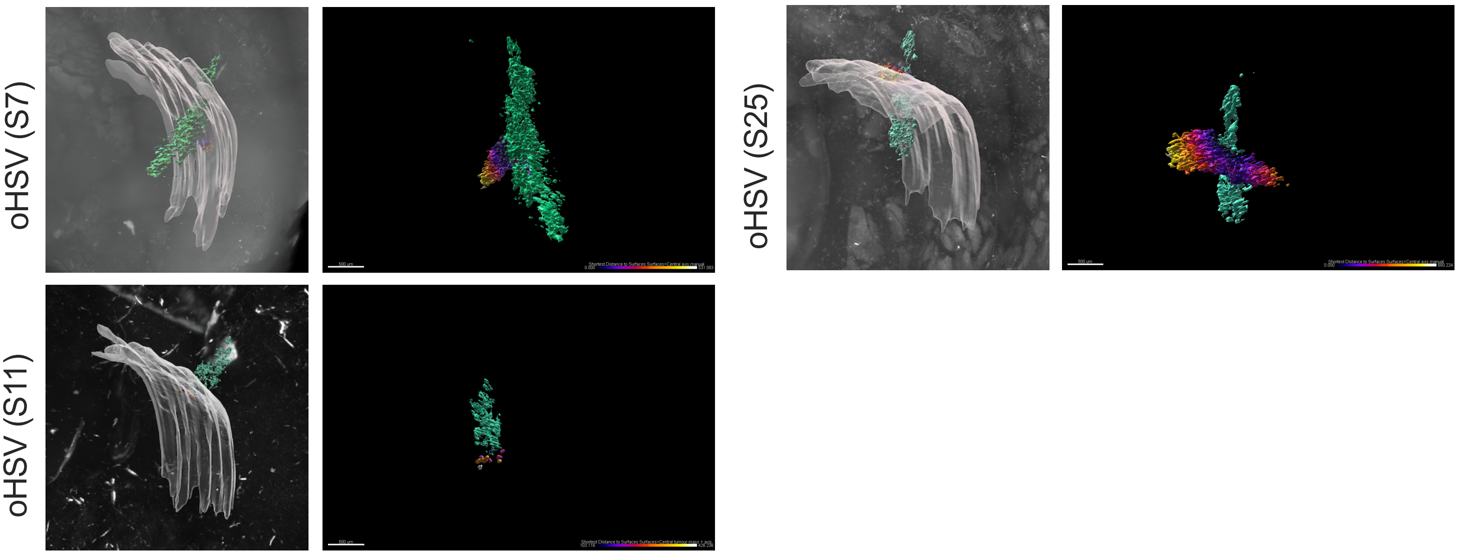


*In vivo* experiment #2

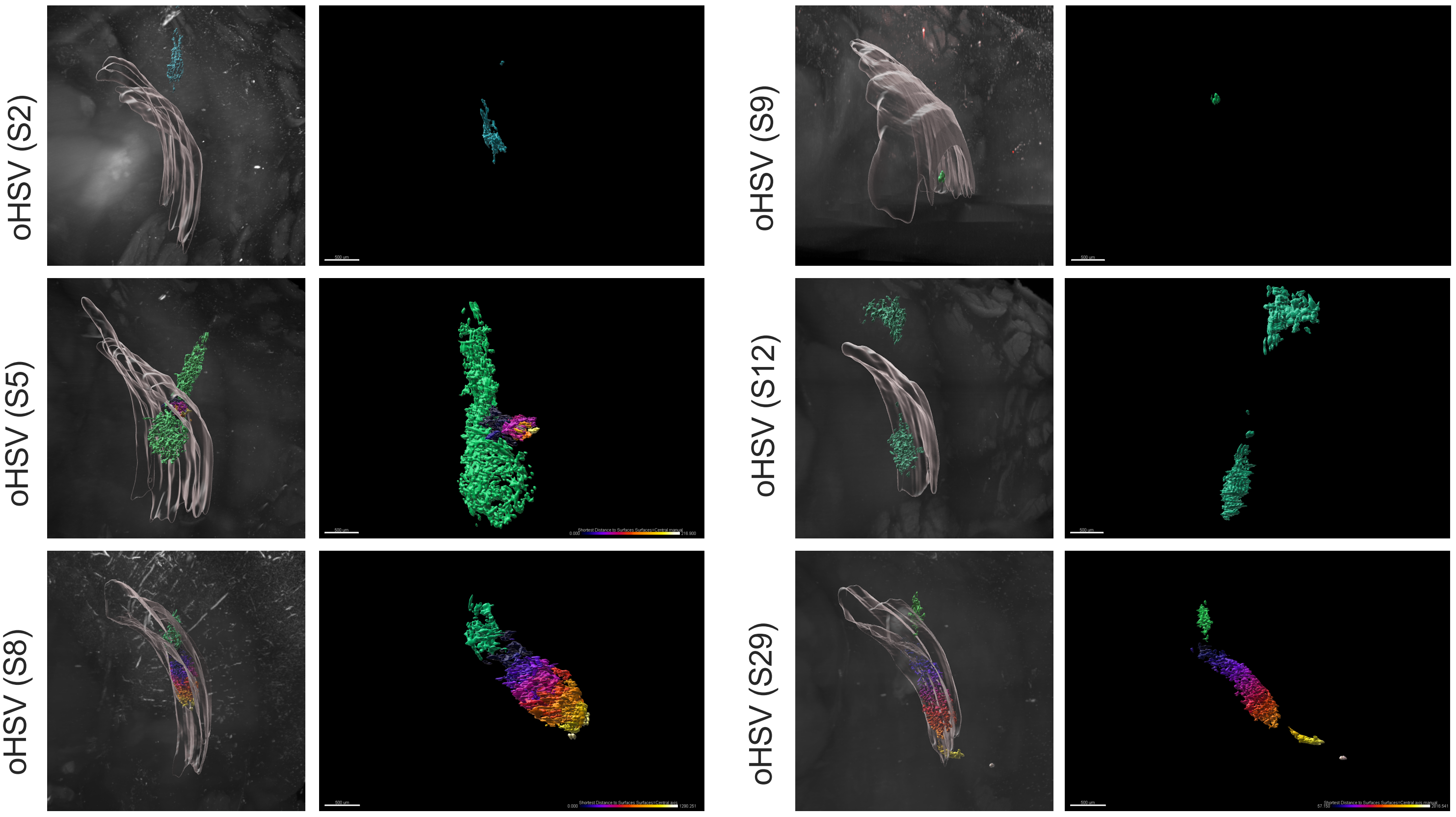

Figure S4-C

*In vivo* experiment #1

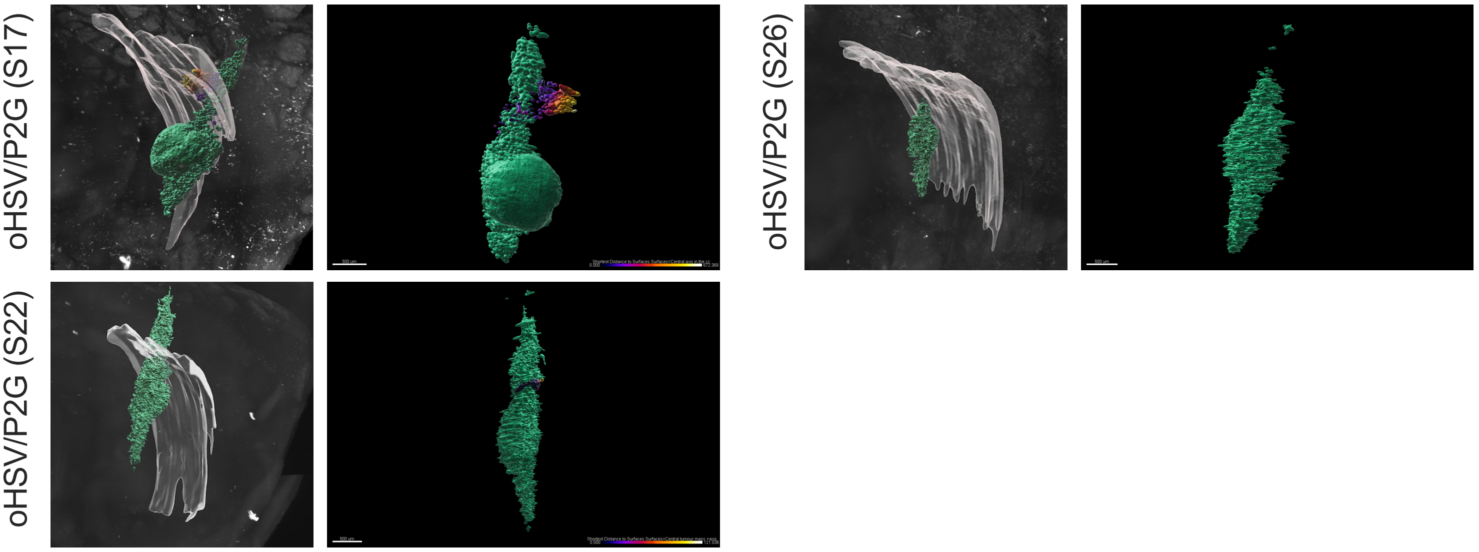


*In vivo* experiment #2

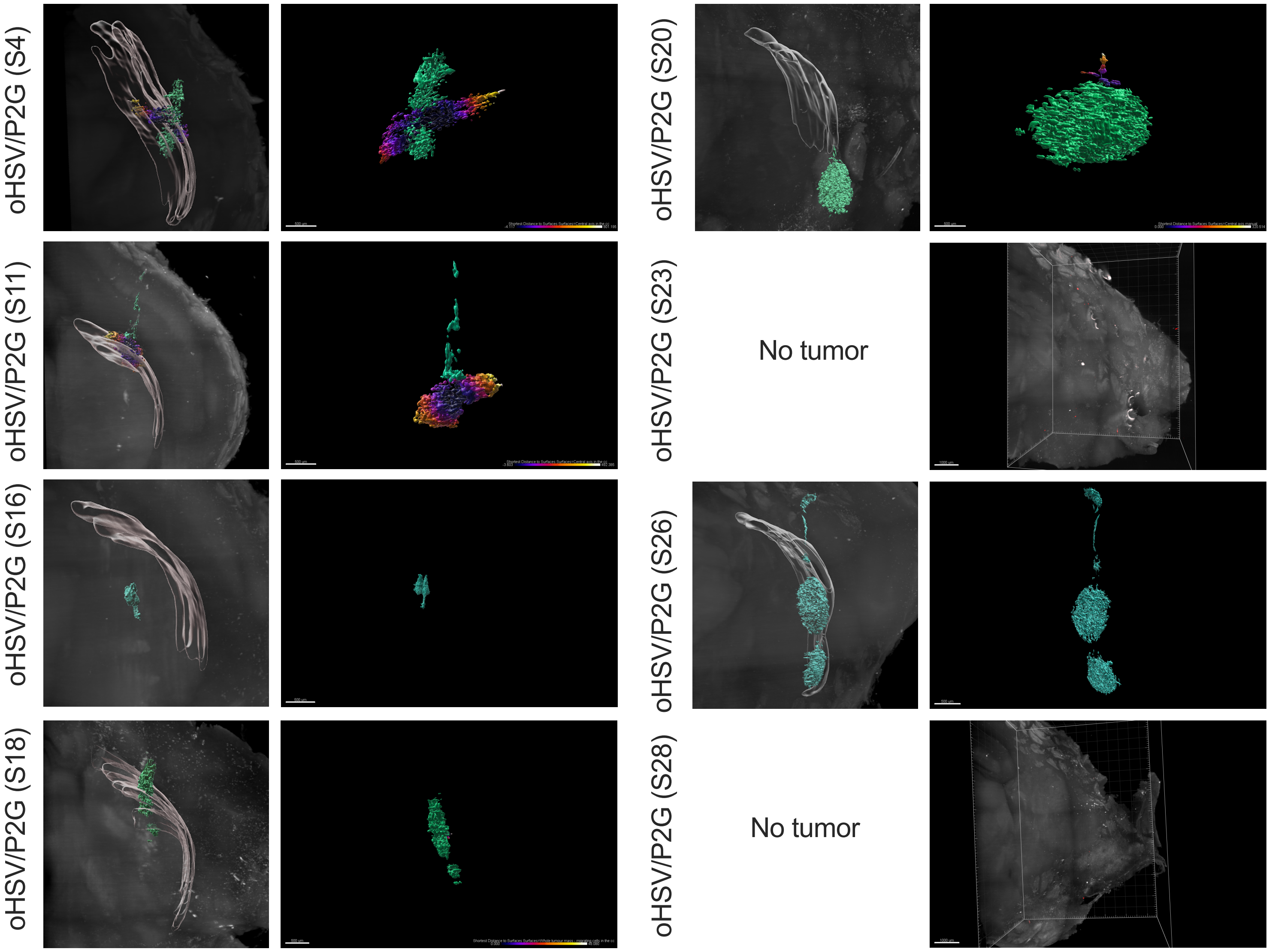


**Figure S4**

Imaris 3D-reconstructions of the tumor mass (green) as well as cells migrating (statistically fire-colored according to their distance to the central tumor mass) through the *corpus callosum* (grey, in the insert) in all the brains analyzed with lighsheet microscopy. Bar represents 500 µm. Quantifications are given in Figures 5C to G.
