## Supplemental material: R script for "An oncolytic herpesvirus expressing a CXCR4 antagonist interferes with Glioblastoma cells stemness features and migration"

**Supplemental material: R script**
**In vivo Exp 1 and 2 (Figure 5)**

#### In vivo Exp 1 and 2: Figure 5B: statistical analysis (R)

```
> # Xenogen signal
> boxcox(data=Test, Xenogen~Replicate + Treatment) # log10 advised
> LMXENO<-lm(data=Test, log10(Xenogen)~Replicate + Treatment)
> LMXENO

Call:
lm(formula = log10(Xenogen) ~ Replicate + Treatment, data = Test)

Coefficients:
     (Intercept)    ReplicateMig_2  TreatmentoHSV-WT      TreatmentPBS
         3.56210          -0.80288           0.36557          -0.06594

> summary(LMXENO) #No significant effect but p-value at 0.0605 for effect
of replicate

Call:
lm(formula = log10(Xenogen) ~ Replicate + Treatment, data = Test)

Residuals:
    Min      1Q  Median      3Q     Max
-2.7592 -0.2050  0.3000  0.5804  1.3153

Coefficients:
                 Estimate Std. Error t value Pr(>|t|)
(Intercept)       3.56210    0.42651   8.352 1.06e-08 ***
ReplicateMig_2   -0.80288    0.40835  -1.966   0.0605 .
TreatmentoHSV-WT  0.36557    0.45702   0.800   0.4313
TreatmentPBS     -0.06594    0.45702  -0.144   0.8864
---
Signif. codes:  0 '***' 0.001 '**' 0.01 '*' 0.05 '.' 0.1 ' ' 1

Residual standard error: 1.015 on 25 degrees of freedom
  (35 observations deleted due to missingness)
Multiple R-squared:  0.165,    Adjusted R-squared:  0.06478
F-statistic: 1.646 on 3 and 25 DF,  p-value: 0.204
```

Conclusion:
**> #Repeating analysis with mixed model as the p-value of replicate is cl
ose to significance in above analysis**

```
> boxcox(data=Test, Xenogen~Replicate + Treatment) # log10 advised
> LME_XENO<-lmer(data=Test, log10(Xenogen)~Treatment + (1|Replicate), REML
=F)
> LME_XENO2<-lmer(log10(Xenogen)~(1|Replicate), data = Test, REML=F)
> anova(LME_XENO,LME_XENO2) #No effect of replicate
Data: Test
Models:
LME_XENO2: log10(Xenogen) ~ (1 | Replicate)
LME_XENO: log10(Xenogen) ~ Treatment + (1 | Replicate)
          npar    AIC    BIC  logLik deviance Chisq Df Pr(>Chisq)
LME_XENO2    3 89.615 93.717 -41.807   83.615
LME_XENO     5 92.544 99.380 -41.272   82.544 1.071  2     0.5854
> summary(glht(LME_XENO, lsm(pairwise ~ Treatment), adjust="tukey"))
```

Conclusion:
**#No effect of treatment**

```
         Simultaneous Tests for General Linear Hypotheses
```

```
Fit: lmer(formula = log10(Xenogen) ~ Treatment + (1 | Replicate),
    data = Test, REML = F)

Linear Hypotheses:
                            Estimate Std. Error z value Pr(>|z|)
(oHSV-P2G) - (oHSV-WT) == 0 -0.38987    0.44008  -0.886    0.649
(oHSV-P2G) - PBS == 0        0.04164    0.44008   0.095    0.995
(oHSV-WT) - PBS == 0         0.43151    0.46122   0.936    0.618
(Adjusted p values reported -- single-step method)

> ##
> LM_XENO<-lm(data=Test, log10(Xenogen)~Replicate + Treatment)
> summary(LM_XENO)
```

Conclusion:
**#No effect treatment but almost significative for replicate (0.06 p valu e)**

```
Call:
lm(formula = log10(Xenogen) ~ Replicate + Treatment, data = Test)

Residuals:
    Min      1Q  Median      3Q     Max
-2.7592 -0.2050  0.3000  0.5804  1.3153

Coefficients:
                 Estimate Std. Error t value Pr(>|t|)
(Intercept)       3.56210    0.42651   8.352 1.06e-08 ***
ReplicateMig_2   -0.80288    0.40835  -1.966   0.0605 .
TreatmentoHSV-WT  0.36557    0.45702   0.800   0.4313
TreatmentPBS     -0.06594    0.45702  -0.144   0.8864
---
Signif. codes:  0 ‘***’ 0.001 ‘**’ 0.01 ‘*’ 0.05 ‘.’ 0.1 ‘ ’ 1

Residual standard error: 1.015 on 25 degrees of freedom
  (35 observations deleted due to missingness)
Multiple R-squared:  0.165,   Adjusted R-squared:  0.06478
F-statistic: 1.646 on 3 and 25 DF,  p-value: 0.204

> AOV_LMXENO<-aov(data=Test, log10(Xenogen)~Replicate + Treatment)
> ###Results
> summary(AOV_LMXENO)
```

Conclusion:
**#No effect (very close to being significative for replicate)**

```
 Df Sum Sq Mean Sq F value Pr(>F)
Replicate    1  4.101   4.101   3.979 0.0571 .
Treatment    2  0.991   0.495   0.480 0.6241
Residuals   25 25.771   1.031
---
Signif. codes:  0 ‘***’ 0.001 ‘**’ 0.01 ‘*’ 0.05 ‘.’ 0.1 ‘ ’ 1
35 observations deleted due to missingness
> TukeyHSD(AOV_LMXENO)
```

Conclusion:
**#No effect for treatment**

```
Tukey multiple comparisons of means
    95% family-wise confidence level

Fit: aov(formula = log10(Xenogen) ~ Replicate + Treatment, data = Test)

$Replicate
                 diff       lwr        upr     p adj
Mig_2-Mig_1 -0.812864 -1.652185 0.02645703 0.0570964

$Treatment
```

| | diff | lwr | upr | p adj |
|---|---|---|---|---|
| oHSV-WT-oHSV-P2G | 0.36496887 | -0.7717078 | 1.5016455 | 0.7066082 |
| PBS-oHSV-P2G | -0.06654214 | -1.2032188 | 1.0701345 | 0.9883520 |
| PBS-oHSV-WT | -0.43151101 | -1.6236675 | 0.7606455 | 0.6443391 |

#### In vivo Exp 1 and 2: Figures 5C, F and G: statistical analysis (R)

#### > # Whole tumor volume (Fig. 5C)

```
> boxcox(data=Test, Whole.tumor.volume+1~Replicate + Treatment) # log10 advised
> plot(lm(data=Test, (Whole.tumor.volume+1)~Replicate + Treatment))
> LMVOL<-lm(data=Test, log10(Whole.tumor.volume+1)~Replicate + Treatment)
> LMVOL

Call:
lm(formula = log10(Whole.tumor.volume + 1) ~ Replicate + Treatment,
    data = Test)

Coefficients:
    (Intercept)    ReplicateMig_2  TreatmentoHSV-WT      TreatmentPBS
        7.5071          -1.2556           0.7928            1.6659

> plot(LMVOL) # Fine
> summary(LMVOL)
```

Conclusion:
**#No significant effect of replicate, nor of treatment.**

```
Call:
lm(formula = log10(Whole.tumor.volume + 1) ~ Replicate + Treatment,
    data = Test)

Residuals:
    Min      1Q  Median      3Q     Max
-6.2515 -0.2616  0.5441  0.8862  2.0707

Coefficients:
                 Estimate Std. Error t value Pr(>|t|)
(Intercept)       7.5071     0.8888   8.446 1.19e-08 ***
ReplicateMig_2    -1.2556     0.8554  -1.468    0.155
TreatmentoHSV-WT   0.7928     0.9477   0.837    0.411
TreatmentPBS       1.6659     0.9821   1.696    0.103
---
Signif. codes:  0 ‘***’ 0.001 ‘**’ 0.01 ‘*’ 0.05 ‘.’ 0.1 ‘ ’ 1

Residual standard error: 2.105 on 24 degrees of freedom
Multiple R-squared:  0.1874,    Adjusted R-squared:  0.08587
F-statistic: 1.845 on 3 and 24 DF,  p-value: 0.1659


> #Mixed model here is not justified but we still check as this mixed model method is used for all other analyses.

> boxcox(data=Test, Whole.tumor.volume+1~Replicate + Treatment) # log10 advised
> LME_MWT<-lmer(data=Test, log10(Whole.tumor.volume+1)~Treatment + (1|Replicate), REML=F)
> summary(LME_MMan)
Linear mixed model fit by maximum likelihood  ['lmerMod']
Formula: log10(Migrating.cells.manually.edited.volume + 1) ~ Treatment +
    (1 | Replicate)
   Data: Test

     AIC      BIC   logLik deviance df.resid
   149.5    156.2    -69.7    139.5       23
```

```
Scaled residuals:
    Min      1Q  Median      3Q     Max 
-1.6722 -1.2508  0.2205  0.9301  1.3582 

Random effects:
 Groups    Name        Variance Std.Dev.
 Replicate (Intercept) 0.000    0.000   
 Residual              8.533    2.921   
Number of obs: 28, groups:  Replicate, 2

Fixed effects:
                Estimate Std. Error t value
(Intercept)       3.6538     0.8807   4.149
TreatmentoHSV-WT   1.2309     1.3129   0.938
TreatmentPBS       4.1156     1.3573   3.032

Correlation of Fixed Effects:
            (Intr) THSV-W
TrtmnHSV-WT -0.671       
TreatmntPBS -0.649  0.435
optimizer (nloptwrap) convergence code: 0 (OK)
boundary (singular) fit: see help('isSingular')

> LME_MWT2<-lmer(log10(Whole.tumor.volume+1)~(1|Replicate), data = Test, REML=F)
> anova(LME_MWT,LME_MWT2)
```

Conclusion of mixed model:
**#No effect of replicate, but we still check as this mixed model method is used for all other analyses.**

```
Data: Test
Models:
LME_MWT2: log10(Whole.tumor.volume + 1) ~ (1 | Replicate)
LME_MWT: log10(Whole.tumor.volume + 1) ~ Treatment + (1 | Replicate)
         npar    AIC    BIC  logLik deviance  Chisq Df Pr(>Chisq)
LME_MWT2    3 128.63 132.63 -61.315   122.63                     
LME_MWT     5 129.24 135.90 -59.620   119.24 3.3901  2     0.1836
> summary(glht(LME_MWT, lsm(pairwise ~ Treatment), adjust="tukey"))
```

Conclusion:
**#No effect**

```
	 Simultaneous Tests for General Linear Hypotheses

Fit: lmer(formula = log10(Whole.tumor.volume + 1) ~ Treatment + (1 | 
    Replicate), data = Test, REML = F)

Linear Hypotheses:
                            Estimate Std. Error z value Pr(>|z|)
(oHSV-P2G) - (oHSV-WT) == 0  -0.8689     0.9145  -0.950    0.608
(oHSV-P2G) - PBS == 0       -1.7943     0.9454  -1.898    0.139
(oHSV-WT) - PBS == 0        -0.9254     0.9887  -0.936    0.617
(Adjusted p values reported -- single-step method)

> ##
> LMWT<-lm(data=Test, log10(Whole.tumor.volume+1)~Replicate + Treatment)
> plot(LMWT)
> summary(LMWT) #No effect
```

```
Call:
lm(formula = log10(Whole.tumor.volume + 1) ~ Replicate + Treatment,
    data = Test)

Residuals:
    Min      1Q  Median      3Q     Max
-6.2515 -0.2616  0.5441  0.8862  2.0707

Coefficients:
                 Estimate Std. Error t value Pr(>|t|)
(Intercept)       7.5071     0.8888   8.446 1.19e-08 ***
ReplicateMig_2   -1.2556     0.8554  -1.468    0.155
TreatmentoHSV-WT  0.7928     0.9477   0.837    0.411
TreatmentPBS      1.6659     0.9821   1.696    0.103
---
Signif. codes:  0 '***' 0.001 '**' 0.01 '*' 0.05 '.' 0.1 ' ' 1

Residual standard error: 2.105 on 24 degrees of freedom
Multiple R-squared:  0.1874,    Adjusted R-squared:  0.08587
F-statistic: 1.845 on 3 and 24 DF,  p-value: 0.1659
```

```
> AOV_LMWT<-aov(data=Test, log10(Whole.tumor.volume+1)~Replicate + Treatment)
```

**> ###Results Whole tumour volume**

Conclusion for Whole tumor volume (Figure 5C):
> summary(AOV_LMWT) **#No effect**

```
             Df Sum Sq Mean Sq F value Pr(>F)
Replicate     1  11.74  11.738   2.648  0.117
Treatment     2  12.80   6.400   1.444  0.256
Residuals    24 106.37   4.432
> TukeyHSD(AOV_LMWT) #No effect
  Tukey multiple comparisons of means
    95% family-wise confidence level

Fit: aov(formula = log10(Whole.tumor.volume + 1) ~ Replicate + Treatment, data = Test)

$Replicate
                 diff       lwr       upr     p adj
Mig_2-Mig_1 -1.386378 -3.144624 0.3718686 0.1167146

$Treatment
                       diff        lwr      upr     p adj
oHSV-WT-oHSV-P2G 0.7848804 -1.5781883 3.147949 0.6887416
PBS-oHSV-P2G     1.6524853 -0.7904637 4.095434 0.2297931
PBS-oHSV-WT      0.8676050 -1.6870788 3.422289 0.6773615
```

**> # Migrating cells volume (Based on manually edited 3D pict. (Fig. 5F)**

```
> boxcox(data=Test, Test$Migrating.cells.manually.edited.volume+1~Replicate + Treatment) #log10 a
dvised
> LME_MMan<-lmer(data=Test, log10(Migrating.cells.manually.edited.volume+1)~Treatment + (1|Re
plicate), REML=F)
> summary(LME_MMan)
Linear mixed model fit by maximum likelihood  ['lmerMod']
Formula: log10(Migrating.cells.manually.edited.volume + 1) ~ Treatment +
    (1 | Replicate)
   Data: Test

     AIC      BIC   logLik deviance df.resid
   149.5    156.2    -69.7    139.5       23

Scaled residuals:
    Min      1Q  Median      3Q     Max
-1.6722 -1.2508  0.2205  0.9301  1.3582

Random effects:
 Groups    Name        Variance Std.Dev.
 Replicate (Intercept) 0.000    0.000
 Residual              8.533    2.921
Number of obs: 28, groups:  Replicate, 2

Fixed effects:
                Estimate Std. Error t value
(Intercept)       3.6538     0.8807   4.149
TreatmentoHSV-WT   1.2309     1.3129   0.938
TreatmentPBS       4.1156     1.3573   3.032

Correlation of Fixed Effects:
            (Intr) THSV-W
TrtmnHSV-WT -0.671
TreatmntPBS -0.649  0.435
optimizer (nloptwrap) convergence code: 0 (OK)
boundary (singular) fit: see help('isSingular')

> LME_MMan2<-lmer(log10(Migrating.cells.manually.edited.volume+1)~(1|Replicate), data = Test, R
EML=F)
> anova(LME_MMan,LME_MMan2)
```

Conclusion for migrating cells volume (Figure 5F)
**# Effect of Replicate, so a mixed effect model is justified**

```
Data: Test
Models:
LME_MMan2: log10(Migrating.cells.manually.edited.volume + 1) ~ (1 | Replicate)
LME_MMan: log10(Migrating.cells.manually.edited.volume + 1) ~ Treatment + (1 | Replicate)
          npar    AIC    BIC  logLik deviance Chisq Df Pr(>Chisq)
LME_MMan2    3 153.57 157.57 -73.788   147.57
LME_MMan     5 149.49 156.15 -69.745   139.49 8.086  2    0.01754 *
---
Signif. codes:  0 '***' 0.001 '**' 0.01 '*' 0.05 '.' 0.1 ' ' 1

> summary(glht(LME_MMan, lsm(pairwise ~ Treatment), adjust="tukey"))
```

Conclusion for migrating cells volume (Figure 5F)
**#Effect for P2G compared to PBS**

Simultaneous Tests for General Linear Hypotheses

```
Fit: lmer(formula = log10(Migrating.cells.manually.edited.volume +
    1) ~ Treatment + (1 | Replicate), data = Test, REML = F)

Linear Hypotheses:
                          Estimate Std. Error z value Pr(>|z|)
(oHSV-P2G) - (oHSV-WT) == 0   -1.231      1.313  -0.938  0.61623
(oHSV-P2G) - PBS == 0        -4.116      1.357  -3.032  0.00669 **
(oHSV-WT) - PBS == 0         -2.885      1.419  -2.032  0.10437
---
Signif. codes:  0 '***' 0.001 '**' 0.01 '*' 0.05 '.' 0.1 ' ' 1
(Adjusted p values reported -- single-step method)
> ##
> LMMan<-lm(data=Test, log10(Migrating.cells.manually.edited.volume+1)~Replicate + Treatment)
> plot(LMMan) # OKish
> summary(LMMan)
```

Conclusion for migrating cells volume (Figure 5F)

**# Effect of treatment**

```
Call:
lm(formula = log10(Migrating.cells.manually.edited.volume + 1) ~
    Replicate + Treatment, data = Test)
Residuals:
    Min      1Q  Median      3Q     Max
-4.8933 -3.1889  0.3891  2.1087  4.4324

Coefficients:
                 Estimate Std. Error t value Pr(>|t|)
(Intercept)         4.893      1.282   3.817 0.000836 ***
ReplicateMig_2     -1.704      1.234  -1.381 0.179923
TreatmentoHSV-WT    1.128      1.367   0.825 0.417558
TreatmentPBS        3.941      1.417   2.782 0.010351 *
---
Signif. codes:  0 '***' 0.001 '**' 0.01 '*' 0.05 '.' 0.1 ' ' 1

Residual standard error: 3.037 on 24 degrees of freedom
Multiple R-squared:  0.306,	Adjusted R-squared:  0.2192
F-statistic: 3.527 on 3 and 24 DF,  p-value: 0.03009

> AOV_LMMan<-aov(data=Test, log10(Migrating.cells.manually.edited.volume+1)~Replicate + Treat
ment)
```

**> ###Results MMan**

```
> summary(AOV_LMMan) # Effect of treatment, not of replicate
            Df Sum Sq Mean Sq F value Pr(>F)
Replicate    1  24.45   24.45   2.651 0.1165
Treatment    2  73.14   36.57   3.966 0.0325 *
Residuals   24 221.32    9.22
---
Signif. codes:  0 '***' 0.001 '**' 0.01 '*' 0.05 '.' 0.1 ' ' 1
> TukeyHSD(AOV_LMMan)
```

Final Conclusion for migrating cells volume (Figure 5F)

#Effect for P2G compared to PBS.

```
  Tukey multiple comparisons of means
    95% family-wise confidence level
```

```
Fit: aov(formula = log10(Migrating.cells.manually.edited.volume + 1) ~ Replicate + Treatment, data =
Test)
$Replicate
                diff       lwr       upr     p adj
Mig_2-Mig_1 -2.000711 -4.536843 0.535422 0.1165464

$Treatment
                      diff        lwr      upr     p adj
oHSV-WT-oHSV-P2G 1.109617 -2.2989242 4.518159 0.6987623
PBS-oHSV-P2G     3.910967  0.3872046 7.434730 0.0275966 *
PBS-oHSV-WT      2.801350 -0.8835812 6.486281 0.160859
```

**> # Migrating cells volume ratio (Based on manually edited 3D pict.) (Fig. 5G)**

```
> boxcox(data=Test, Test$Migrating.cells.manually.edited.volume.ratio+0.0001~Replicate + Treatmen
t) # log10 advised
> LME_MMan_r<-lmer(data=Test, log10(Migrating.cells.manually.edited.volume.ratio+0.0001)~Treat
ment + (1|Replicate), REML=F)
> summary(LME_MMan_r)
Linear mixed model fit by maximum likelihood  ['lmerMod']
Formula: log10(Migrating.cells.manually.edited.volume.ratio + 1e-04) ~
    Treatment + (1 | Replicate)
   Data: Test

     AIC      BIC   logLik deviance df.resid
   104.2    110.9    -47.1     94.2       23

Scaled residuals:
    Min      1Q  Median      3Q     Max
-1.7849 -1.0423  0.2627  0.5714  1.9341

Random effects:
 Groups    Name        Variance Std.Dev.
 Replicate (Intercept) 0.000    0.000
 Residual              1.694    1.302
Number of obs: 28, groups:  Replicate, 2

Fixed effects:
                 Estimate Std. Error t value
(Intercept)       -2.6434     0.3924  -6.736
TreatmentoHSV-WT   0.9665     0.5850   1.652
TreatmentPBS       2.0248     0.6048   3.348

Correlation of Fixed Effects:
            (Intr) THSV-W
TrtmnHSV-WT -0.671
TreatmntPBS -0.649  0.435
optimizer (nloptwrap) convergence code: 0 (OK)
boundary (singular) fit: see help('isSingular')

> LME_MMan2_r<-lmer(log10(Migrating.cells.manually.edited.volume.ratio+0.0001)~(1|Replicate), da
ta = Test, REML=F)
> anova(LME_MMan_r,LME_MMan2_r)
```

Conclusion
### Effect of Replicate, so a mixed effect model is justified

```
Data: Test
Models:
LME_MMan2_r: log10(Migrating.cells.manually.edited.volume.ratio + 1e-04) ~ (1 | Replicate)
LME_MMan_r: log10(Migrating.cells.manually.edited.volume.ratio + 1e-04) ~ Treatment + (1 | Replica
te)
            npar    AIC    BIC logLik deviance  Chisq Df Pr(>Chisq)
LME_MMan2_r    3 109.68 113.68 -51.84  103.681
LME_MMan_r     5 104.22 110.88 -47.11   94.219 9.4616  2    0.00882 **
---
Signif. codes:  0 ‘***’ 0.001 ‘**’ 0.01 ‘*’ 0.05 ‘.’ 0.1 ‘ ’ 1
> summary(glht(LME_MMan_r, lsm(pairwise ~ Treatment), adjust="tukey"))
```

Conclusion (Figure 5G):
**#Effect for P2G compared to PBS.**

```
                Simultaneous Tests for General Linear Hypotheses

Fit: lmer(formula = log10(Migrating.cells.manually.edited.volume.ratio +
    1e-04) ~ Treatment + (1 | Replicate), data = Test, REML = F)

Linear Hypotheses:
                          Estimate Std. Error z value Pr(>|z|)
(oHSV-P2G) - (oHSV-WT) == 0  -0.9665     0.5850  -1.652  0.22374
(oHSV-P2G) - PBS == 0       -2.0248     0.6048  -3.348  0.00237 **
(oHSV-WT) - PBS == 0        -1.0584     0.6324  -1.674  0.21514
---
Signif. codes:  0 '***' 0.001 '**' 0.01 '*' 0.05 '.' 0.1 ' ' 1
(Adjusted p values reported -- single-step method)

> ##
> LMMan_r<-lm(data=Test, log10(Migrating.cells.manually.edited.volume.ratio+0.0001)~Replicate + T
reatment)
> plot(LMMan_r) # OKish
> summary(LMMan_r)
```

Conclusion (Figure 5G)
**# effect of treatment**

```
Call:
lm(formula = log10(Migrating.cells.manually.edited.volume.ratio +
    1e-04) ~ Replicate + Treatment, data = Test)

Residuals:
    Min      1Q  Median      3Q     Max
-2.1487 -1.2139  0.3328  0.6050  2.6600

Coefficients:
                Estimate Std. Error t value Pr(>|t|)
(Intercept)       -2.2629     0.5830  -3.881 0.000711 ***
ReplicateMig_2    -0.5232     0.5612  -0.932 0.360448
TreatmentoHSV-WT   0.9347     0.6217   1.504 0.145723
TreatmentPBS       1.9713     0.6443   3.060 0.005382 **
---
Signif. codes:  0 '***' 0.001 '**' 0.01 '*' 0.05 '.' 0.1 ' ' 1

Residual standard error: 1.381 on 24 degrees of freedom
Multiple R-squared:  0.3117,    Adjusted R-squared:  0.2256
F-statistic: 3.622 on 3 and 24 DF,  p-value: 0.02747

> AOV_LMMan_r<-aov(data=Test, log10(Migrating.cells.manually.edited.volume.ratio+0.0001)~Repli
cate + Treatment)



> ###Results MMan_r
> summary(AOV_LMMan_r) # Effect of treatment, not of replicate
            Df Sum Sq Mean Sq F value Pr(>F)
Replicate    1   2.81   2.807   1.472  0.237
Treatment    2  17.92   8.960   4.698  0.019 *
Residuals   24  45.77   1.907
---
Signif. codes:  0 '***' 0.001 '**' 0.01 '*' 0.05 '.' 0.1 ' ' 1
> TukeyHSD(AOV_LMMan_r) #Effect for P2G compared to PBS.
  Tukey multiple comparisons of means
```

```
  95% family-wise confidence level

Fit: aov(formula = log10(Migrating.cells.manually.edited.volume.ratio + 1e-04) ~ Replicate + Treatmen
t, data = Test)

$Replicate
                  diff       lwr       upr     p adj
Mig_2-Mig_1 -0.6779209 -1.831303 0.4754611 0.2368993

$Treatment
                      diff        lwr      upr     p adj
oHSV-WT-oHSV-P2G 0.9253659 -0.6247700 2.475502 0.3129546
PBS-oHSV-P2G     1.9555024  0.3529662 3.558039 0.0147678 *
PBS-oHSV-WT      1.0301365 -0.6456959 2.705969 0.2928678
```
